# Local B cell maturation and mast cell regulation of choroid plexus function in early life.

**DOI:** 10.1101/2025.11.27.690721

**Authors:** Samir Ali-Moussa, Ana Blas-Medina, Joseph M Josephides, Olivier Mirabeau, Inès Bouteau, Tamara Matijevic, Mariangeles Kovacs, Victoria Pakulska, Andrei Reyes Pangan, Christian Thomas, Martin Hasselblatt, Xinzhong Dong, Mariette Matondo, Nicolas Gaudenzio, Friederike Jönsson, Laetitia Travier, Aleksandra Deczkowska

## Abstract

Postnatal development is a critical period for the maturation of the nervous and immune systems. The choroid plexus (CP) within the brain ventricles guides brain development through the production of cerebrospinal fluid and responds to stimuli from its local immune microenvironment. Here, using single-cell sequencing, we chart the establishment of the immune niche within the CP from birth to adulthood. We demonstrate that the CP is an active site for the development of B cells from early pro-B cells to mature B cells. We also characterize a transient population of CP mast cells that is highly abundant in the perinatal period. Single activation of these cells shortly after birth led to activation of serotonin-dependent secretion from the CP epithelial cells and resulted in cognitive impairment later in life. Our findings highlight the crucial nature of the CP as a neuroimmune interface, where cellular crosstalk regulates key functions of CP activity, thereby guiding brain development.

## Introduction

Postnatal development is a critical window for brain maturation, during which foundational neural circuits are established and refined in response to both intrinsic genetic programs and extrinsic environmental cues(*1, 2*). Characterized by intense neurological rewiring, including synaptogenesis, gliogenesis, myelination, and activity-dependent synaptic pruning(*3, 4*), this phase also coincides with key events in immune system maturation, including the development of bone marrow hematopoiesis, the emergence of adaptive immunity and dynamic changes in immune cell phenotypes in response to external stimuli, such as the forming microbiota(*5, 6*). The neuroimmune communication processes that take place within this window are critical for the healthy maturation of the brain, and their perturbation in paradigms of maternal and postnatal immune activation were shown to lead to atypical brain development, resulting in altered behavioral outcomes in adulthood(*7–9*). In humans, early-life infections, allergies, microbiome dysbiosis, and inborn errors of immunity have been linked to the manifestation of neurodevelopmental disorders such as intellectual delays, attention deficit and hyperactivity disorder, schizophrenia, and autism spectrum disorder(*10–13*).

Yet, immune signals have limited access to the brain, since their presence is anatomically restricted to the brain borders, such as the meningeal layers, the circumventricular organs, and the choroid plexus (CP)(*14–17*). The CP, a specialized structure positioned within the ventricles and responsible for the production of cerebrospinal fluid (CSF), plays an essential role in shaping healthy brain development by enriching the CSF in key hormones, signaling proteins, and growth factors through diverse secretory mechanisms(*18–22*). This CP secretory activity is critical during development, regulating various developmental processes, from neuroprogenitor proliferation and differentiation to the survival of various cell types within the brain(*23, 24*). At the same time, the local immune cells that populate the CP stroma(*16*), as well as the expression of cytokine and toll-like receptors by the CP epithelium(*14*), make it an active neuroimmune niche that relays signals from the periphery to the brain, across various physiological and pathological conditions(*20, 25–31*).

In this study, we use single-cell RNA sequencing to map the immune landscape of the CP from birth to adulthood, revealing the heterogeneity and dynamics of local immune populations. Using this approach, we found that the postnatal CP harbors a diversity of developing B cells, from progenitor through cycling pre-B cells to mature B cells, which persists into adulthood. We further describe a population of CP mast cells (MCs) that inhabit the CP transiently at birth. Using pharmacological and chemogenetic approaches, we show that a single activation of CP MCs (CPMCs) at birth results in a serotonin-mediated activation of secretion from the CP epithelium, which leads to cognitive deficits later in life. These results provide possible explanation of the link between MC-triggering infections, such as meningitis and early life allergies, and persistent intellectual impairment(*32, 33*). Together, our results provide fundamental understanding as to how changes in local immune function within the CP during postnatal development may regulate its secretory properties and shape brain development through the CSF.

## Results

### The immune landscape of the CP across postnatal development

To comprehensively map the immune cell heterogeneity and dynamics in the postnatal developing CP, we micro-dissected CPs from mice at different timepoints from birth to adulthood: postnatal day 0 (P0; birth), P6 (immature CP and gut epithelial barriers)(*30*), P10 and P15 (early juvenile mice), P21 (weaning period when microbiome drives maturation of immune system)(*34*), and P49-63 adults (Ad), and performed Cellular Indexing of Transcriptomes and Epitopes by sequencing (CITE-seq) for simultaneous measurement of transcriptomic and surface protein expression on FACS-sorted CD45^+^ immune cells (Fig. 1A, and fig. S1A and B). Unsupervised clustering and dimensionality reduction were performed on a total of 30414 cells (4803 from P0, 4671 from P6, 4631 from P10, 4319 from P15, 5352 from P21, and 6637 from Ad) with a weighted nearest neighbor analysis using both RNA and surface protein expression simultaneously (Fig. 1B and fig. S2, A and B). We obtained a total of 31 clusters aggregated onto 16 main lineages, including macrophages: Macs1 (*Csf1r^+^, Adgre1^+^, C1qa^+^*), Macs2 (*Zeb2^+^, Fcgr3^+^*), proliferating macrophages (*Mki67^+^, Top2a^+^*), and epiplexus cells (*Cd9^+^, Spp1^+^, Mrc1^-^*), monocytes (*Ly6c^+^, Ccr2^+^*), conventional type one and type two dendritic cells : cDC1 (*Flt3^+^, Zbtb46^+^, Xcr1^+^*), and cDC2 (*Flt3^+^, Zbtb46^+^, Cd209a^+^),* respectively, B cells (*Cd19^+^, Pax5^+^*), T cells (*Cd3e^+^, Lat^+^*), NK cells (*Cd3e^-^, Xcl1^+^, Klrb1c^+^*) and other lymphocytes, including innate lymphoid cells and gamma delta T cells (*Gata3^+^, Cd163l1^+^*), neutrophils (*Ly6g^+^, S100a9^+^*), MCs (*Kit^+^,Cpa3^+^, Gata2^+^, Fcer1a^+^*), and basophils (*Kit^-^, Gata2^+^, Fcer1a^+^*), as well as non-immune cells of epithelial (*Ttr^+^, Ecrg4^+^*), mesenchymal (*Pdgfra^+^,Lum^+^*), and endothelial (*Plvap^+^, Cd34^+^*) lineages. The cellular identities were further validated at the surface protein level through key surface markers (Fig. 1, C and D, and fig. S2C).

**Fig. 1.**
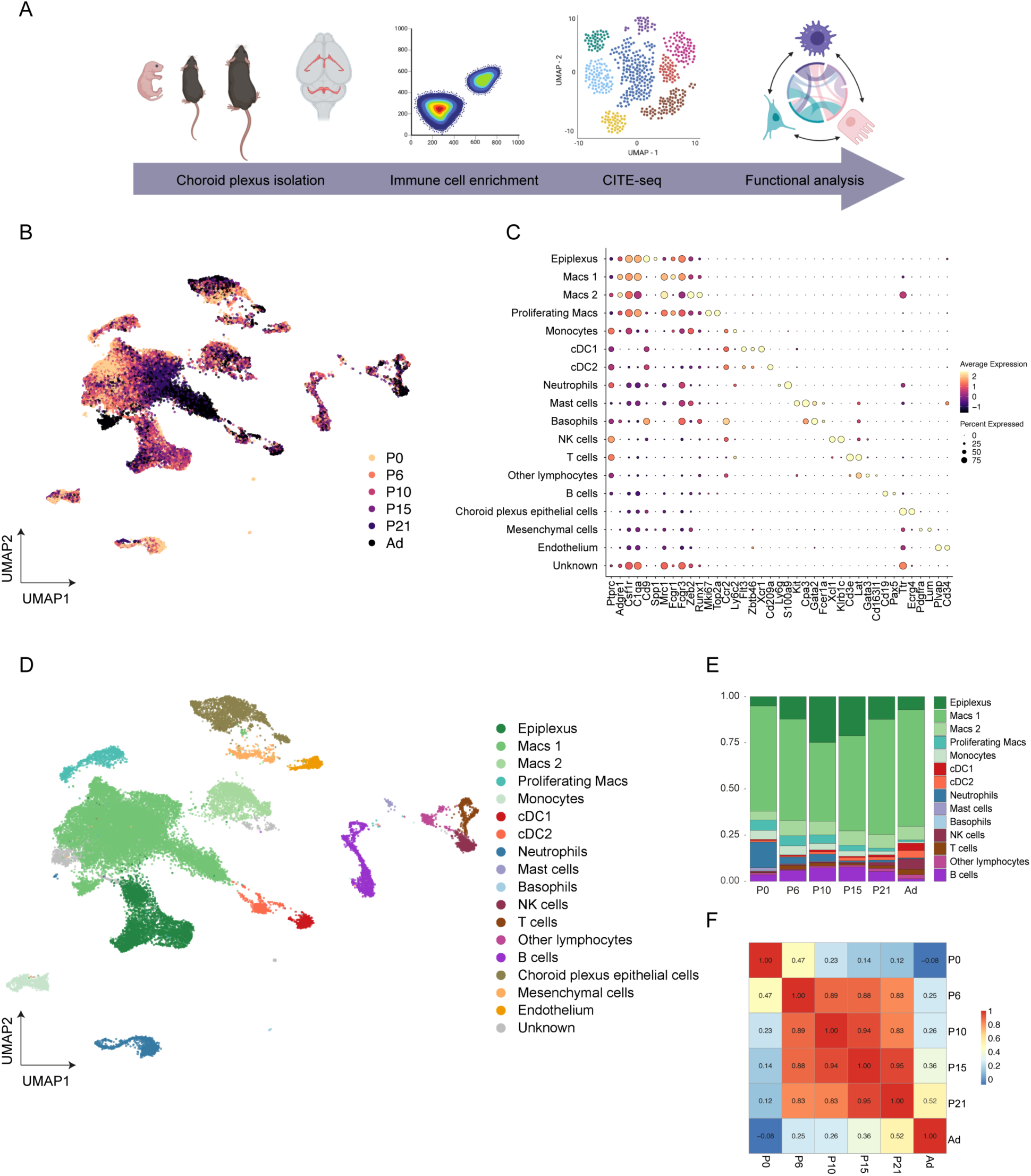
Mapping immune cell dynamics during postnatal development in the CP. **(A)** Schematic illustration of the CITE-seq pipeline. **(B)** Integrated Uniform Manifold Approximation and Projection (UMAP) visualization of the CITE-seq data across postnatal development (30414 cells; 4803 from P0, 4671 from P6, 4631 from P10, 4319 from P15, 5352 from P21, and 6637 from Ad). **(C)** Dot plot showing normalized expression of selected marker genes for the main cell types. The color represents mean expression level, and the size indicates the proportions of cells expressing the genes. **(D)** UMAP visualization of the identified cell types in the CP. **(E)** Bar plots showing the immune composition of the CP during postnatal development (subsampled on 3000 immune cells per age). **(F)** Heatmap displaying Pearson correlation values between cluster proportions across pairs of samples, from P0 to Ad.

The immune landscape of the CP underwent dynamic changes across postnatal development; granulocytes, including neutrophils, MCs and basophils, as well as proliferating macrophages were more abundant at P0 and decreased gradually over time. B cells and epiplexus cells gradually increased, peaking at P15, then waning until adulthood, while cDC1, cDC2, and non-B lymphocytes gradually increased throughout, peaking in adulthood (Fig. 1E). By examining the correlation in cluster proportion across developmental stages, we found that P0 and Ad clustered distinctly, whereas P6, P10, P15 and P21 were more similar, reflecting the presence of a dynamic maturation process in immune cell composition during postnatal development, and suggesting that the CP immune ecosystem undergoes its most significant changes during the period when CP function is critical for brain maturation(*35–38*) (Fig. 1F, and fig. S2, D and E).

### The CP is a site of B cell development

Our analysis revealed an enrichment of B cells in the CP at P15, followed by a progressive decline until adulthood (Fig. 1E). This transient surge of CP B cells suggests a temporally regulated expansion or infiltration event during this window. This interesting pattern prompted us to further investigate CP B cells. We first confirmed this progression using flow cytometry, which demonstrated that live CD45^+^ CD19^+^ B220^+^ B cells increased from P0 to P15 and were significantly less abundant in adult mice (Fig. 2A and fig. S3A). Using immunohistochemistry, we further validated the presence of B220^+^ B cells within the P15 CP stroma (Fig. 2B).

**Fig. 2.**
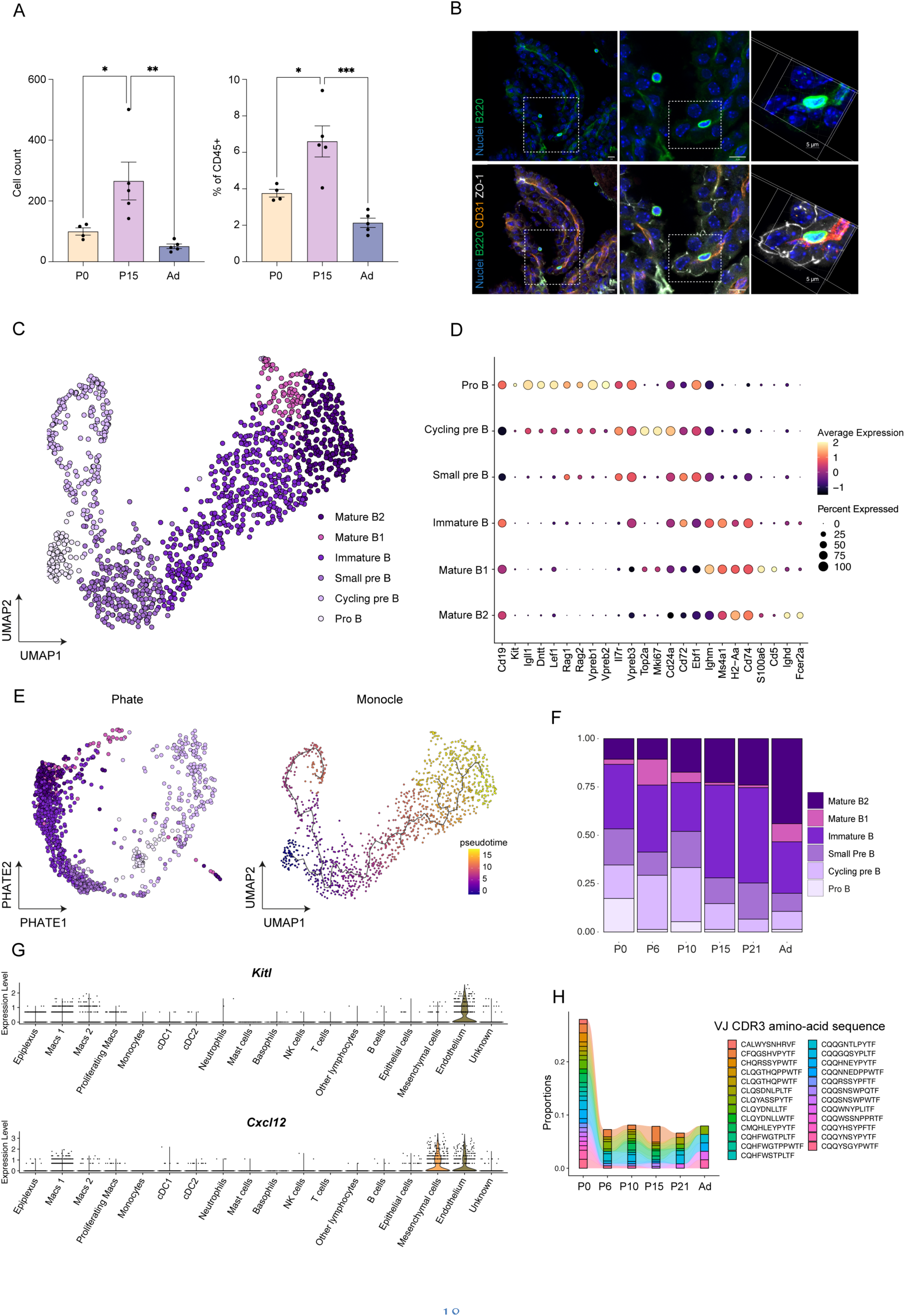
The CP is a developmental niche for B cell development. **(A)** Flow cytometry quantification of CD19^+^ B220^+^ B cell numbers (left) and percentages (right). n= 4-5 mice per group. Data are represented as mean ± SEM, one-way ANOVA with Tukey’s multiple comparison test. **(B)** Representative confocal image of 0.8µm-thick CP section (left and middle panels) and three-dimensional reconstruction (right panels) of the same but 10μm-thick CP imaged area highlighting B cells (B220), vasculature (CD31), epithelial cell junctions (ZO-1) and nuclei (DAPI). 10µm (left and middle panels, 5µm (right panel). **(C)** UMAP visualization of subclustered B cell subpopulations. **(D)** Dot plot showing normalized expression of selected marker genes for B cell subpopulations. The color represents mean expression level, and the size indicates the proportions of cells expressing the genes. **(E)** Trajectory inference analysis (PHATE and Monocle3) confirms the progression from pro-B cells to mature B cells in the CP. **(F)** Bar plots showing the proportion of the different B cell subpopulations across postnatal development. **(G)** Violin plot depicting the expression of *Cxcl12* and *Kitl* across all cell types. **(H)** Amino-acid composition of the Variable-Junctional segment of the hypervariable CDR3 across postnatal development. *p < 0.05; **p < 0.01; ***p < 0.001.

Interestingly, by subclustering B cells separately, we were able to detect all developmental stages of B cells in the postnatal CP, including pro B cells (*Kit^+^, Dntt^+^, Rag 1^+^, Rag2^+^*), cycling pre B cells (*Mki67*^+^, *Top2a*^+^), small pre B cells (*Ebf1^+^*, *Ighm^+^)*, immature (*Ms4a1^+^*, *H2-Aa^+^),* mature B1 cells (*Cd5^+^, S100a6^+^*) as well as mature B2 cells (*Ighd^+^, Fcer2a^+^).* This annotation was also reflected by the expression of key surface markers (Fig. 2, C and D, and fig. S3B), suggesting that, similarly to the meninges, the CP is a site of local B cell maturation(*39*). Pseudotime trajectory analysis and PHATE confirmed the progression from progenitor to mature B cells, usually observed in the bone marrow(*39, 40*) (Fig. 2E, and fig. S3C). Along development, B cell progenitors were more abundant in early postnatal stages, peaking at P15, while mature B cell frequencies were highest in adult mice (Fig. 2F). We further confirmed this distribution through flow cytometry using markers for mature B2 cells (IgD^+^ CD23^+^), mature B1 cells (IgD^-^, CD23^-^, CD5^+^), immature B cells (IgM^+^), pro B cells (CD43^+^), pre B cells (CD43^-^) (fig. S3, A and D).

In the bone marrow and meninges, B cell development was shown to be guided by essential niche factors provided by stromal cells(*39*). Similarly, in our dataset, the CP mesenchymal and endothelial cells expressed *Cxcl12* and *Kitl*, which encodes the Stem Cell Factor, suggesting that the CP stroma may act as a support to guide postnatal development of progenitor B cells in the CP (Fig. 2G).

Given that B cell development requires a rigorous step of selection in which autoreactive B cells are eliminated(*40*), we next questioned whether such a process took place *in situ* in the postnatal CP by analyzing B cell clonality through B cell receptor sequencing (BCR-seq). The BCR compartment in the CP was composed mainly of IgM and IgD expressing B cells whereas other immunoglobulin heavy chain subclasses were absent (Fig. 3F). This suggests that, in contrast to the meninges(*39, 41*), B cell activation and class switching do not spontaneously occur in the CP. Additionally, the BCR repertoire analysis showed that the complementarity-determining region 3 (CDR3), a high variability region, displayed higher richness at P10 and P15, at the peak of B cell development in the CP, and decreased in adulthood (fig. S3F). Furthermore, the amino-acid sequences of the CDR3 region demonstrated a highly diverse repertoire at P0 that progressively decreased over postnatal development, culminating in the persistence of only four B cell clones in the adult CP (Fig. 2H).

**Fig. 3.**
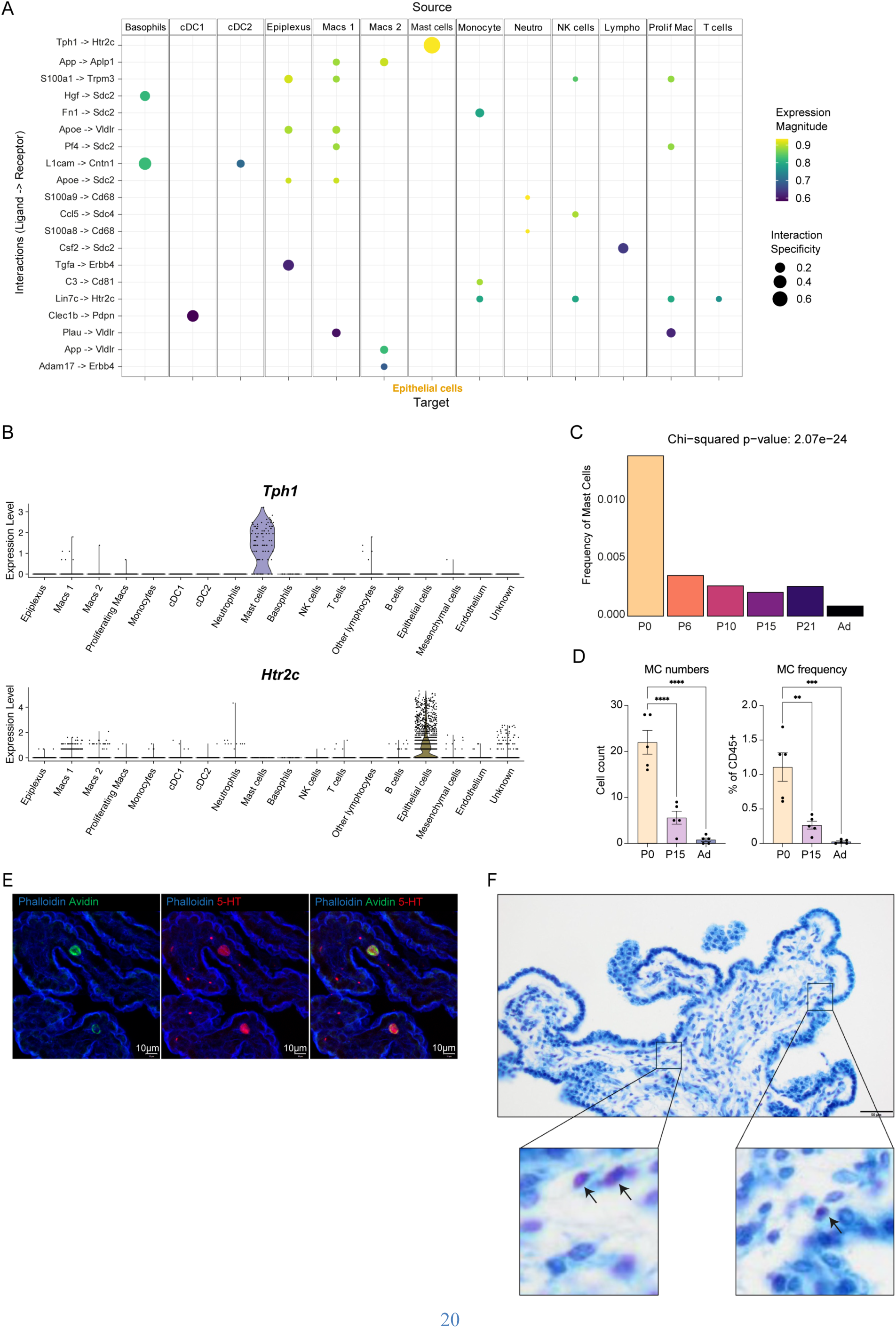
Neonatal CPMCs interact with epithelial cells through serotonin. **(A)** Dot plot of cell-cell communication between immune cells and CPECs. Each dot represents ligand-receptor pairs. Dot size represents the specificity assigned by LIANA, while the color represents the magnitude of the interaction. **(B)** Violin plots depicting the expression of *Tph1* and *Htr2c* across all cell types. **(C)** Bar plot showing the proportion of MCs in the CP across all ages in the CITE-seq data. **(D)** Flow cytometry plots (upper panel) and quantification (lower panel) of Lineage^-^ c-KIT^+^ CD200R^+^ MCs numbers (left) and percentages (right). n= 5 mice per group. Data are represented as mean ± SEM, one-way ANOVA with Tukey’s multiple comparison test. **(E)** Representative confocal image and 10.4µm-thick maximum intensity projection of 4V CP sections highlighting MCs (Avidin), serotonin (5-HT), epithelium and vasculature (Phalloidin). Bar: 10µm. **(F)** CP from a P0 donor stained with toluidine blue. Purple cells (arrows in insets) represent granule-filled MCs. Bar: 50 µm. **p < 0.01; ***p < 0.001; ****p < 0.0001.

Together, our findings suggest that B cell maturation and clonal selection occur *in situ* within the CP leading to the establishment of a restricted and persistent B cell repertoire.

### CPMCs interact with epithelial cells through serotonin

We next sought to identify the pathways of immune-CP communication, which could potentially alter CP function and thereby impact postnatal brain development. To this end, we performed ligand-receptor inference analysis between CP epithelial cells (CPECs) and immune cells across all ages. This analysis revealed a potent and specific interaction between tryptophan hydroxylase 1 (*Tph1*) encoding for the enzyme catalyzing the conversion of tryptophan into serotonin, expressed by MCs, with the 5-hydroxytryptamine receptor 2C (*Htr2c*), encoding for the serotonin receptor, expressed by CPECs (Fig. 3A). The expression of both *Tph1* and *Htr2c* was restricted to MCs and CPECs, respectively (Fig. 3B). In the brain, *Htr2c* was highly expressed by the CP in development (fig. S4A).

The CP epithelium plays a key role in the maintenance of the electrochemical balance of the CSF through diverse active and passive transport systems, while facilitating the clearance of waste metabolites produced as a result of neuronal activity(*23, 35*). Beyond that, the CP epithelium produces a repertoire of morphogens and hormones, including Wnt and Sonic-hedgehog signaling proteins, Insulin and Insulin-like growth factors that directly act on neighboring central nervous system (CNS)-residing cells to shape their phenotypes and functions(*37*). Building on recent findings that demonstrated a critical role for the HTR2C receptor in the regulation of CP secretory activity during embryonic period(*22*), we sought to determine whether local MCs contribute to the modulation of CP secretory function in postnatal development.

To start, we set off to characterize the CPMCs. Analysis of cell proportion in our CITE-seq dataset showed that MCs were more abundant in the neonatal CP at P0 and were rarely found in other timepoints (Fig. 3C). To verify this quantitatively, we analyzed MCs using flow cytometry with previously established markers; MCs were gated on live, lineage-negative, hematopoietic cells that concomitantly express CD200R and c-KIT as per previous reports (CD45^+^, CD11b^-^, CD3^-^, TER119^-^, Ly6G^-^, CD19^-^, NK1.1^-^, CD200R^+^, C- KIT^+^)(*42*). Indeed, the number and percentage of CPMCs decreased gradually from P0, when MCs represented about 1% of immune cells, to adults, when they were virtually absent (Fig. 3D). Our analysis is consistent with that of a previous transcriptomic atlas of the CP (Dani, et al. 2021(*19*)), in which the presence of *Tph1*-expressing MC was restricted to the E16.5 CP, representing 1% of immune cells, while being absent in both the adult and aged mouse CP (fig. S4, B and C). Imaging of MCs using avidin(*43*) confirmed the presence of serotonin-positive MCs(*43*) in the stroma of the mouse CP at P0 (Fig. 3E). Furthermore, using toluidine staining(*44*), we detected MCs in a postmortem neonatal human CP tissue, suggesting that their presence within the early postnatal CP is conserved between mice and humans (Fig. 3F).

MCs can be classified into two main subtypes: connective tissue MCs (CTMCs), found in the skin and peritoneal cavity, and mucosal MCs (MMCs), located in mucosal tissues such as the gut and lung(*45*). To better understand CPMCs biology, we integrated our CPMC data with a published single-cell dataset of MMCs and CTMCs (Tauber et al(*45*)) as well as dural MCs (Rustenhoven et al(*17*)), which were recently shown to be of a CTMC phenotype (fig. S4D). Using previously established markers(*45*) (Table S1), we scored cells from each tissue based on their CTMC, or MMC phenotype. We were able to determine that CPMCs, displayed a high CTMC signature score, associated with high expression of core CTMC genes such as *Mrgprb2, Cpa3, Cd81, Mcpt4,* and *Cma1*, suggesting a connective tissue phenotype (fig S4, E and F). We next used flow cytometry to validate this identity and probe for markers of MC maturation in CPMCs at P0(*42*). We used Mrgprb2-Cre-GFP mouse line in which GFP is expressed under the promoter of the CTMC-specific receptor Mrgprb2(*46*). All CPMCs were positive for GFP, similarly to MCs in the skin and dura at this timepoint. Further, across the three organs, MCs displayed low expression of the IgE receptor FceR1a, and high expression of the IL33 receptor ST2, suggesting that CPMCs may have limited capacity for IgE-mediated activation, which is consistent with prior reports demonstrating that MCs mature postnatally by upregulating the FceR1a(*42*) (fig. S4G). Notably, CPMCs also displayed higher expression of *Tph1* when compared to other tissue MCs signatures, suggesting that CPMCs may be uniquely poised as serotonin-producing cells in the postnatal brain (fig. S4H). Taken together, our results uncover a novel transient population of MCs within the neonatal CP capable of serotonin production.

### CPMC degranulation induces CPEC activation

We next set out to explore the relationship between CPMCs and CPECs in neonates. Recent work has shown that serotonin triggers apocrine secretion of CPECs in both embryos and adult, via activation of the HTR2C receptor(*20, 22*). We therefore hypothesized that MC activation may change CPEC secretion through serotonergic signaling.

To set off, we used compound 48/80 (C48/80), a known agonist of the CTMC-specific receptor Mrgprb2(*47*). To target CNS-residing MCs while minimizing off-target activation of peripheral MCs, we administered C48/80, or saline as a control, intracerebroventricularly (i.c.v) at birth. MC activation through C48/80 induced a potent recruitment of neutrophils, monocytes, and NK cells to the CP 24h post injection, with no changes in Macs, T cells, B cells or MC numbers (fig. S5A), similarly to previously reported consequences of MC activation in other tissues(*48–50*).

To assess the involvement of serotonergic signaling in MC-dependent CP activation, neonates also received 1μg of the selective HTR2C antagonist SB242084 (SB), or saline (Sal), injected intraperitoneally (i.p) one hour prior to C48/80 administration. We quantified CPEC apocrine events, a hallmark of serotonergic signaling activation in the CP(*20, 22*), three hours after C48/80 administration (Fig. 4A). To do so, we stained for apical surface markers of the CP epithelium, aquaporin 1 (AQP1) and Phalloidin, as well as zonula-1 (ZO-1), to delineate the boundaries of individual CPECs, and quantified the cells with marked apical membrane discontinuities. C48/80-dependent CP activation induced a two-fold increase in the number of apocrine secretory events by CPECs, at both the LV and 4^th^ ventricle (4V) CP. This effect was inhibited by the HTR2C antagonist SB (Fig. 4, B and C, and fig. S5B), indicating that MC-derived serotonin regulates CPEC secretion.

**Fig. 4.**
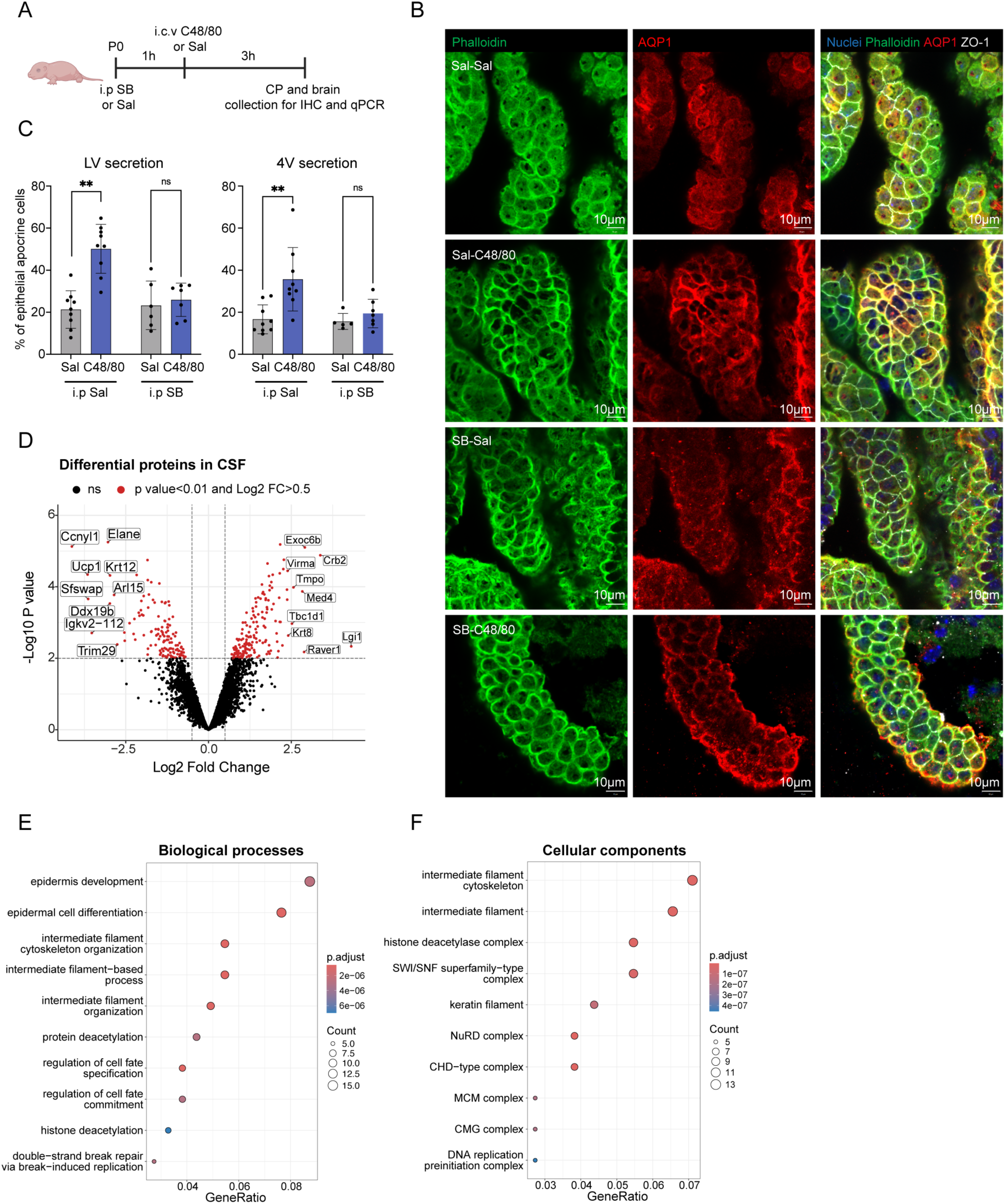
Neonatal MC activation induces CPEC activation and apocrine secretion. **(A)** Experimental design. **(B)** Representative confocal imaging of the top 0.8 µm-thick surface of CPECs from 4V CP sections of P0 mice, three hours following injection with C48/80 or Sal, with and without prior inhibition with SB. Sections were stained for nuclei, Phalloidin, AQP1 and ZO-1. Bar: 10µm. **(C)** Quantification of apocrine CPECs from **(B)** in the 4V (right panel) and also in the LV CP (left panel), three hours following MCs activation with C48/80, with and without SB (n= 5-9 mice per group). **(D)** Volcano plot of the differentially expressed proteins in the CSF of C48/80 and saline injected mice. **(E)** Dot plot showing top10 statistically most significant BP terms enriched in the C48/80 CSF proteome. **(F)** Dot plot showing top10 statistically most significant CC terms enriched in the C48/80 CSF proteome. **p < 0.01; ns: p>0.05.

Given the observed effects of MC activation on CPEC secretory activity, we next sought to assess the effects of these changes on CSF proteomic composition. To this end, CSF was collected from the cisterna magna of neonatal mice injected with either saline or the C48/80, followed by mass spectrometry-based proteomic analysis. A total of 242 proteins were upregulated in the CSF of C48/80 injected mice (Fig. 4D). To better interpret the biological significance of these changes, we performed gene ontology (GO) analysis for biological processes (BP) and cellular component (CC). Enriched BP terms included proteins involved in epidermis development, cytoskeleton organization and cell fate commitment, processes typically associated with intracellular structural remodeling and differentiation (Fig. 4E). CC analysis further revealed a predominance of proteins originating from intracellular compartments, including the cytoskeleton and the nucleus, such as histone deacetylase complex, ATPase complex, and the minichromosomal maintenance (MCM) complex (Fig. 4F). These findings suggest that MC activation induces a significant upregulation of apocrine secretion, resulting in the release of intracellular CPECs’ content into the CSF.

CSF composition can reflect neuronal activity and broader CNS responses. To parse CP-specific effect of MC activation from general CNS responses, we cultured P0 CP explants in artificial CSF (aCSF), stimulated with C48/80 (or Sal) for 3h, and analyzed the aCSF proteome. Using this method, we identified a total of 1080 proteins significantly upregulated in the aCSF of C48/80 stimulated explants (fig. S5C). GO BP analysis revealed an upregulation of terms involved in mRNA processing, catabolic processes, and regulation of translation (fig. S5D). CC analysis showed a predominance of proteins belonging primarily to the nuclear speck and the ribosomal subunits (fig. S5E), further supporting the hypothesis MC activation drives the release of CPEC intracellular content onto the CSF. Together, these data show that MC-derived serotonin induces CPEC apocrine secretion, potentially altering CSF composition with CPEC intracellular content and shaping brain maturation during postnatal development.

### Perinatal activation of CPMCs or CP secretion leads to cognitive deficits in adulthood

MC activation is a classic hallmark of allergic inflammation, and MC activation syndrome is commonly associated with neurodevelopmental disorders, altered behavior and cognitive impairment(*33*). Beyond allergies, adult dural MCs have recently been shown to act as first responders in bacterial meningitis(*50*), an infection typically occurring in neonates, where they can be directly activated through the Mrgprb2 receptor(*48*). Importantly, neonatal meningitis has also been associated with permanent focal deficits and cognitive impairment(*32*).

To investigate the long-lasting consequences of perinatal MC activation on brain development, we activated MCs at birth with a single i.c.v injection of C48/80, allowed the mice to age to eight weeks and performed a battery of behavioral tests (Fig. 5A). Transient MC activation did not impact baseline anxiety (elevated plus maze, the open field), locomotor activity (open field) nor social preference (three-chamber test) (fig. S6, A-C). Strikingly, C48/80 mice exhibited impaired recognition memory in the novel object recognition test, with no changes in total interaction time suggesting that transient MC activation within the brain at birth is sufficient to induce long-lasting cognitive deficits later in life (Fig. 5B).

**Fig. 5.**
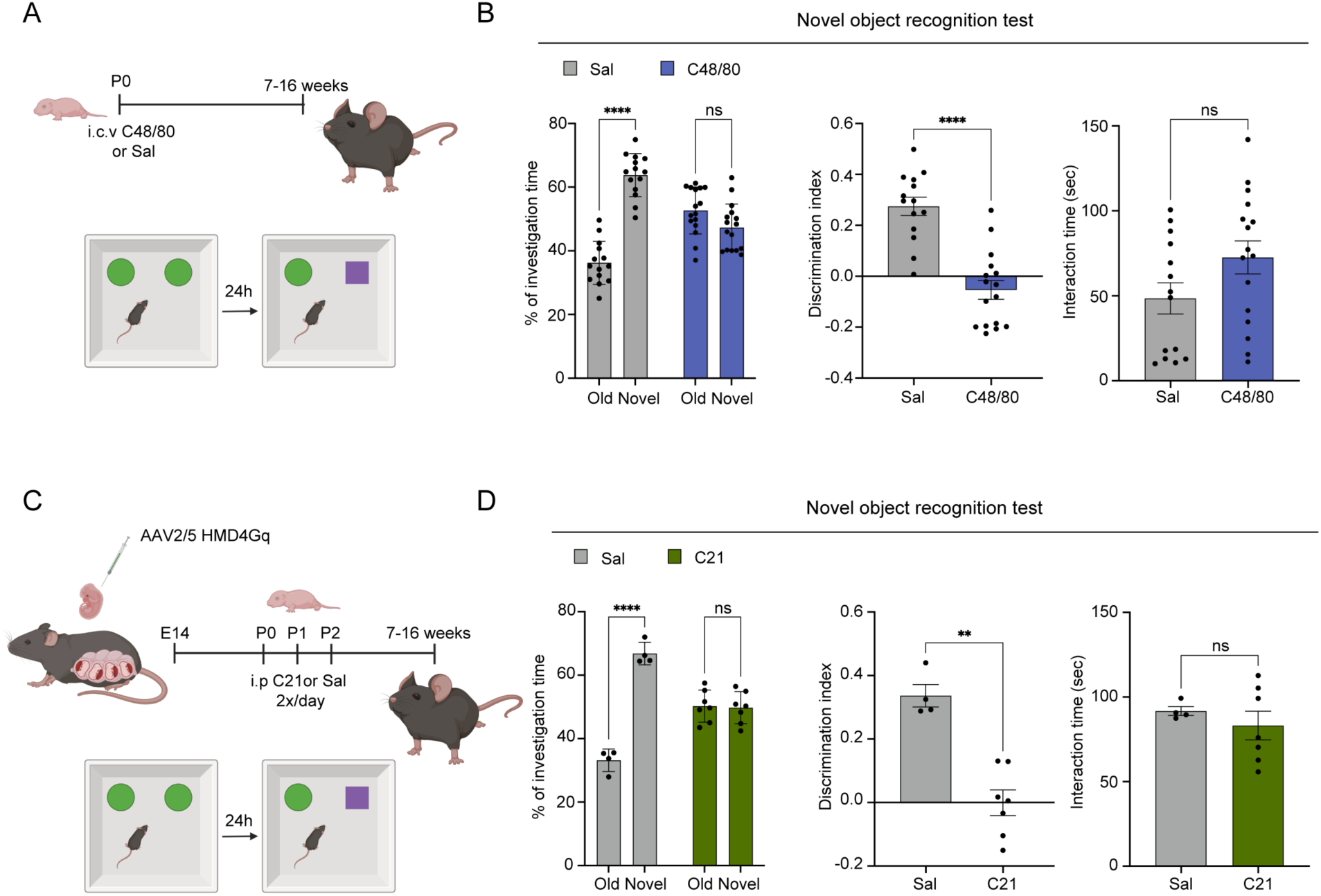
Neonatal MC activation and chemogenetic CPEC activation induce long-lasting cognitive deficits epithelial. **(A)** Experimental design. **(B) Left**: Percentage of interaction time spent with the novel versus old object for control and C48/80-injected mice (*n* = 14-16 mice per group from two independent experiments). Data were analyzed by two-way ANOVA with Tukey’s multiple comparisons test. **Center**: Discrimination index for control and C48/80 mice (calculated as the difference in interaction time between the novel and old object over total interaction time). Data were analyzed by Mann-Whitney test. **Right**: total interaction time during the novel object interaction test by control and C48/80 mice. Data were analyzed by Mann-Whitney test. **(C)** Experimental design. **(D) Left**: Percentage of interaction time spent with the novel versus old object for saline and C21-injected DREADD mice (*n* = 4-7 mice per group from one experiment). Data were analyzed by two-way ANOVA with Tukey’s multiple comparisons test. **Center**: Discrimination index for saline and C21-injected DREADD mice. Data were analyzed by Mann-Whitney test. **Right**: total interaction time during the novel object interaction test by saline and C21-injected DREADD mice. Data were analyzed by Mann-Whitney test. **p < 0.01; ****p < 0.0001; ns: p>0.05.

MCs were previously found in various areas of perinatal brain(*44*). To isolate the functional effect of transient MC-mediated CPEC activation on the developing brain, independently of MC-induced neuroinflammation, we developed a chemogenetic strategy to selectively activate the CP *in vivo* within a restricted perinatal time-window. Given that HTR2C is a Gq protein-coupled receptor, we created a system for CP-specific expression of the excitatory designer receptor exclusively activated by designer drugs (HMD4Gq). To deliver the HMD4Gq construct specifically to the CP, we packed it into the CP-tropic adeno-associated virus (AAV) serotype AAV2/5(*26, 51*) and injected it i.c.v on embryonic day 14 (E14) (fig. S6D).

This approach enabled the targeted overexpression of the receptor at birth, which we validated using immunostainings, demonstrating the specificity and transduction efficiency of AAV2/5 (fig. S6E). Upon i.p administration of compound 21 (C21), the selective agonist of the HMD4Gq receptor, we observed a marked increase in the expression of the immediate early gene c-FOS in CPECs, confirming the functional efficacy of our approach (fig. S6F). We next assessed whether abnormal activation of CPECs is responsible for the observed behavioral and cognitive deficits. To do so, we activated CPECs using C21, with saline serving as a control (injected twice a day, for 3 days to account for the short half-life of C21, and the long-lasting effect of MC activation(*52*), and performed behavioral tests on the same mice in adulthood (Fig. 5C). Similarly to the MC activation model, perinatal activation of CPEC resulted in similar cognitive deficits measured on the novel object recognition test when compared to their saline injected littermates (Fig. 5D), with no differences observed in other behavioral metrics (fig. S6, G-I). These cognitive deficits were also detected in the novel object location test (fig. S6J), suggesting spatial memory deficits. Thus, both MC activation and CPEC activation in the neonatal period resulted in long-lasting cognitive impairments, suggesting that abnormal MC activation may drive these cognitive deficits through the modulation of CPEC secretory activity.

## Discussion

Peripheral signals of immune and microbial origin have been shown to shape tissue maturation from embryogenesis to postnatal development, and in virtually all organs(*53–55*). Within the CNS, immune cells at the brain borders and within the parenchyma have been demonstrated to promote key events of brain development(*14, 53, 56–58*). For example, meningeal innate lymphoid cells have been shown to guide inhibitory cortical neuron development postnatally(*58*), while parenchymal MCs and T cells have been reported as key drivers of male sexual maturation and microglia development, respectively(*59, 60*). Previous studies have consistently focused on the meningeal layers and parenchyma-resident immune cells. Here, we suggest that the CP provides another route for immune regulation of brain development.

We show that the CP harbors a diverse immune population undergoing an extensive transformation during the postnatal period. Our data tracking B cell phenotypes and BCR over time strongly suggest that B cells mature locally within the CP during postnatal development, in a process that closely resembles that of B cell development in the bone marrow and in the meninges(*39, 40*). However, B cell development in the CP was more active during the neonatal period, with only a few progenitors remaining in adult mice, suggesting that the CP retains its function as a niche for B cell development later in life, albeit at a lower scale. The origin and function of CP B cells is yet to be addressed, and so is their antigen specificity.

We also characterize a transient population of CPMCs that are present in the CP early in postnatal development. Recent studies have positioned MCs as key regulators of CSF production and flow in pathological conditions. At the CP, MCs were shown to increase CSF production through tryptase release, which disturbs the CPEC cilia in a mouse model of tumor-associated hydrocephalus(*61*). In the meninges, dural MCs were shown trigger histamine-induced vasodilation of bridging veins at the arachnoid cuff exits points and thereby modulate CSF efflux from the brain territory(*50*). Our findings expand the current paradigms by which MCs regulate CSF properties; using the selective agonist of the Mrgprb2 receptor, we demonstrate that early life MC activation within the CNS, induces apocrine secretion of CPEC, potentially altering CSF protein composition.

Importantly, we show that early-life MC activation and associated CP secretion lead to persistent cognitive deficits later in life. These findings align with clinical observations linking neonatal meningitis and mastocytosis to long-term cognitive impairments(*32, 33*). We propose that MCs are strategically positioned at the CP during the neonatal period, where the blood-CSF barrier is immature and neonates are highly susceptibility to meningitis(*30*). In this context MCs serve a protective role as a first line of defense against pathogens, orchestrating the immune response(*50*). However, as a result of their activation, MC-induced serotonergic signals alter the CP’s secretory function during a critical window of brain development, resulting in long-lasting cognitive deficits. This model has broad implications for understanding how immune signals at brain interfaces contribute to neurodevelopmental disorders and opens new avenues for therapeutic intervention targeting MCs or CP secretory pathways to mitigate neurodevelopmental disorders linked to early-life inflammation.

## Supplementary

### Materials and Methods

#### Animal procedures

Wild-type 8–10 weeks old and neonatal (P0–P21) C57BL/6J/RJ mice were bred in-house or purchased from the Janvier labs. MrgprB2-EGFP/Cre mice were provided by Xinzhong Dong.

All mouse strains were maintained at Institut Pasteur, in a specific pathogen–free facility, and housed in a 12-hour light-dark cycle with water and rodent chow *ad libitum*. All the procedures were in agreement with the guidelines of the European Commission for the handling of laboratory animals, directive 86/609/EEC, approved by the ethical committees No. 59, by the CETEA/CEEA No. 089, under the number dap210067 and APAFIS #32382-2021070917055505.

#### Human samples

The use of archived human samples for research upon anonymization was in accordance with local regulations of the University Hospital Münster and approved by the Münster ethics committee (2007-420-f-S and 2017-707-f-S).

#### *In utero* intracerebroventricular injection of adeno-associated viruses

Timed pregnant females at E14.5 were anesthetized with isoflurane, injected subcutaneously with buprenorphine, and placed on a heat-pad. A laparotomy was performed and 1μl of AAV2/5-CMV::hM3Dq GFP (VectorBuilder) at a titer of 1×10^10^ viral particles/ul was introduced into the lateral ventricle of each embryo. The dams were then sutured and monitored until delivery. Foster dams were used from birth to weaning. Chemogenetic activation of newborn pups was performed by an intraperitoneal injection of compound 21 (1 mg/kg).

#### CSF sampling of pups

Mice were anesthetized and the neck was disinfected with ethanol. A glass capillary was inserted into the cisterna magna to collect the CSF, which was then transferred to protein-low-binding tubes and frozen at - 80°C until further analysis. CSF from 3-5 mice was pooled per sample for proteomic analysis.

#### Artificial CSF preparation and explant culture

aCSF was freshly prepared on the day of the experiment with concentrations closely mimicking physiological neonatal CSF(*35*). The following final concentrations in mM were used: Na⁺ 116, Cl⁻ 109, K⁺ 10, Ca²⁺ 4, Mg²⁺ 1, HCO₃⁻ 26, and H₂PO₄⁻ 1 (as NaH₂PO₄). The CPs from all four ventricles were dissected and placed in a 96 well plate with 50ul of aCSF. CP explants were stimulated 50uM of C48/80 for 3h at 37°C and 5% CO₂. Following stimulation, the aCSF was carefully collected and frozen at -80°C until analysis.

#### Liquid chromatography - Mass spectrometry LC-MS/MS analysis

CSF, and aCSF samples were processed using the iST Sample Preparation Kit (PreOmics GmbH, Germany) following the manufacturer’s instructions, with the following modification. Proteins were digested overnight at 37 °C using mass-spectrometry–grade Trypsin/Lys-C (Promega, #V507A) at a 50:1 protein-to-enzyme ratio in 100 mM triethylammonium bicarbonate (TEAB). Resulting peptides were resuspended in 2% acetonitrile and 0.1% formic acid. Peptide concentration was determined by measuring absorbance at 280 nm using a NanoDrop 2000 spectrophotometer (Thermo Scientific), and an amount equivalent to 25 ng of peptides was injected for LC-MS/MS analysis.

Peptide mixtures were analyzed on a NanoElute 2 UHPLC system (Bruker Daltonics) coupled to a timsTOF Ultra 2 mass spectrometer (Bruker Daltonics) operated in DIA-PASEF mode. Peptides were separated on a PepSep Ultra C18 analytical column (25 cm × 75 μm, 1.5 μm; Bruker Daltonics) maintained at 50 °C. Chromatographic separation was performed at a flow rate of 0.25 µL/min using buffer A (0.1% formic acid in water) and buffer B (0.1% formic acid in acetonitrile). The 30-min gradient consisted of: 2% to 4% B in 1 min, 4% to 20% B from 1 to 20 min, 20% to 25% B from 20 to 25 min, followed by an increase to 95% B in 1 min and a hold at 95% B for 4 min.

The timsTOF Ultra 2 was operated in DIA-PASEF acquisition mode using 24 isolation windows of 25 m/z each, covering a precursor mass range of 400–1000 m/z. Instrument parameters were set as follows: capillary voltage, 1600 V; dry gas flow, 3 L/min; dry temperature, 200 °C. TIMS settings included an ion mobility range of 0.64–1.45 V·s/cm² (1/k₀), with a ramp time of 100 ms and 100% duty cycle.

#### Protein identification and quantification

Raw MS data were processed in Spectronaut (v.20.2.250922.92449; Biognosys) using the directDIA mode with BGS Factory Settings. Searches were performed against the *Mus musculus* UniProt protein database (*54,824 entries*) using the UniProt FASTA parsing rule. The directDIA+ (Deep) workflow with Pulsar Search was used for peptide identification. Carbamidomethylation of cysteine was set as a fixed modification, while protein N-terminal acetylation and methionine oxidation were specified as variable modifications (maximum of five per peptide). Enzyme specificity was set to Trypsin/P and LysC/P, allowing up to two missed cleavages, and peptide length was restricted to 7–52 amino acids.

PSM, peptide, and protein group identifications were controlled at a 1% FDR. Precursor Q-value threshold was 0.01, and protein group Q-value cutoffs were set to 0.01 at the experiment level and 0.05 at the run level. Protein inference was performed using the IDPicker algorithm.

Quantification was carried out at the MS2 level using peak area and the automatic LFQ strategy, selecting the top 1–3 peptides per protein group and 1–3 precursors per peptide. Cross-run normalization was applied using the automatic strategy with local (non-linear) regression for precision iRT calibration. Interference correction was enabled, requiring at least 2 MS1 and 3 MS2 data points. Dynamic extraction windows were used for both the m/z and ion mobility dimensions with a correction factor of 1.

#### Statistical analysis of mass spectrometry data

To identify proteins with differential abundance between conditions, quantified protein intensities exported from Spectronaut were analyzed. Only protein groups containing at least one peptide unique within the FASTA database (“unique peptide”) were retained. In addition, proteins were required to be quantified in at least two biological replicates of at least one of the two conditions being compared.

Proteins lacking intensity values in one condition (i.e., quantified exclusively in the other) were considered present in one condition and absent in the other. These proteins were set aside as differentially abundant by definition and displayed separately in bar plots, ranked by their iBAQ (intensity-based absolute quantification) values [ https://doi.org/10.1038/nature10098]. For proteins quantified in both conditions, intensity values were log₂-transformed and normalized by median centering within each condition (section 3.5 in [https://doi.org/10.1007/978-1-0716-1967-4_12]).

Differential abundance was assessed using a moderated *t*-test implemented in the **limma** R package [https://doi.org/10.1093/nar/gkv007; https://www.biorxiv.org/content/10.1101/2020.05.29.122770v1.abstract. Resulting *p*-values were corrected for multiple testing using an adaptive Benjamini–Hochberg procedure via the *adjust.p* function of the **cp4p** package [https://doi.org/10.1002/pmic.201500189, employing the robust estimation method described in [https://doi.org/10.1093/bioinformatics/btl328 o infer the proportion of true null hypotheses. Proteins identified as statistically significant by this analysis, together with proteins present in only one condition, were reported as differentially abundant and visualized in volcano plots. ClusterProfiler was used to perform GO analysis on differentially upregulated proteins.

#### Tissue collection and Flow cytometry

Mice were euthanized with an intraperitoneal injection of a lethal dose of anesthetics (150mg/kg of ketamine and 25mg/kg of xylazine) and transcardially perfused with Phosphate Buffer Saline (PBS). The CP, dura mater, tongue, skin, and brain tissues were harvested and collected in Roswell Park Memorial Institute medium (RPMI) 1640 medium and mechanically dissociated using fine scissors (dura, CP, tongue, skin) and a homogenizer (brain). For the CP, dura, skin, and tongue, an additional enzymatic dissociation with a mix of 10 U/ml collagenase I, 400 U/ml collagenase IV and 30 U/ml DNase I diluted in RPMI was performed for 30 minutes at 37°C under mild agitation, followed by mechanical disruption by pipetting. After dissociation, the cells were washed with FACS buffer (0.5% Bovine Serum Albumin (BSA), 0.4% Ethylenediaminetetraacetic acid (EDTA), in PBS+/+), filtered through a 70 µm cell strainer, and centrifuged at 400 g for 5 min at 4°C to obtain a single cell suspension. The cells were then resuspended in blocking buffer (1:100 anti-mouse CD16/CD32 in FACS buffer) to prevent unspecific epitope labeling and incubated 10 minutes on ice. Antibodies against cell surface epitopes (**Table S2)** were then added for 30 min in the dark followed by a viability dye for an additional 30 min. The cells were then washed, resuspended in FACS buffer, and acquired using the Sony ID7000. Data were analyzed using FlowJo software (BD Biosciences).

#### Tissue collection for immunostaining

Mice were euthanized as above and perfused first with PBS+/+ (to maintain the integrity of the CP epithelial cell junctions), followed by 4% paraformaldehyde (PFA). The brains were collected, post-fixed in PFA 4% for 90 minutes, washed in PBS for 16-48 hours at 4 °C, and embedded in 30% sucrose, 0.5% sodium azide in PBS at 4 °C until fully saturated for cryopreservation. Brains were cut along the midline and the two hemispheres were embedded in Optimal Cutting Temperature (OCT) compound, solidified on dry ice, and stored at -80 °C. Sagittal cryosections were cut using a cryostat (Leica), directly mounted onto glass slides, and stored at -20 °C until further processing.

Slides were thawed at room temperature for 10 minutes and rinsed in PBS^-/-^ to remove OCT. Slides were dried and a hydrophobic barrier was drawn using a hydrophobic pen. Sections were blocked and permeabilized in PBST (PBS^-/-^ with 0.2% Triton X-100) with 10% normal donkey serum (NDS), and 10% normal goat serum (NGS) (blocking buffer) for 2 hours at RT. Primary antibodies were diluted in blocking buffer and incubated at 4 °C overnight or up to 72 hours for c-Fos staining. After three washes in PBS^-/-^, secondary antibodies were diluted in blocking buffer and incubated 2 hours at RT in the dark. Slides were washed three times in PBS^-/-^, mounted with Fluoromount, covered with a coverslip and dried overnight, and stored at –20 °C. Images were acquired using a confocal Zeiss LSM700 upright microscope.

#### Image analysis

Each image is representative for at least five analyzed animals per condition/age, both male and female, for which several images have been taken and processed.

Apocrine secretion events at the surface of the CPECs was quantified using the plugin “Cell Counter” of the Fiji software: first, the CPECs for which the boundaries were complete were selected in the ZO-1 channel, then, among these selected CPECs, those showing marked apical membrane discontinuities besides cilia in the Phalloidin channel were enumerated.

Three-dimensional reconstructions were performed using ImarisViewer software using planal clipping.

#### Toluidine blue staining

Formalin-fixed, paraffin-embedded tissue sections were deparaffinized in xylene and rehydrated through a graded ethanol series to distilled water. The sections were then stained for 3 minutes with a Toluidine Blue O solution containing 1% Toluidine Blue, 1% disodium tetraborate, and 1% Pyronin.

#### Single cell RNA sequencing

Mice were anaesthetized and perfused as per above. CP tissues were microdissected from mice at P0, P6, and P10 (n=20 per timepoint), P15 and P21 (n=18 each), and adult (n=16 in two replicates) and processed following a previously published protocol(*16*) (with modifications). Specifically, tissues were placed in the caps of DNA lobind Eppendorf tubes with a small volume of ice-cold RPMI containing 30 µM ActD (Sigma, No. A1410) and cut into small pieces with microscissors. The CPs from each mouse was processed separately. The enzyme mix containing 10 U/ml collagenase I, 400 U/ml collagenase IV, 30 U/ml DNase I and 15 µM ActD (diluted in RPMI) was added to the tissues and incubated at 11 °C for 50 min. Every 10 min the solution was gently resuspended. Cells from all samples were washed with MACS buffer with 3 µM ActD. The CPs were then resuspended in 1ml of MACS Buffer 3 µM ActD and pooled together. Blocking was performed with the TruStain FcX^TM^ PLUS for 10min on ice. The cells were then incubated with FITC- anti CD45 for 5 min, followed by the Total-seq antibody cocktail (resuspended according to the manufacturer’s instructions) for 25 min, followed by DAPI for an additional 5 min. After washing, approximately 15,000 cells per sample were sorted on a BD FACS Aria III and loaded into individual wells of a Chromium Chip K (10X Genomics). Libraries were generated using the Chromium next GEM single cell 5’ feature barcode kit (10X Genomics) according to the manufacturer’s instructions. Sequencing was performed on the Nextseq 2000 Illumina sequencing platform.

#### Single-cell RNA analysis

Sequencing reads were processed with Cell Ranger v7.2.0 and aligned to the mm10-2020-A genome. Antibody-derived tag (ADT) counts for CITE-seq were obtained using the TotalSeq-C Mouse Universal Cocktail (BioLegend; Antibody Reference 199903). Lymphocyte clonotype analysis was performed using the Cell Ranger V(D)J mouse reference refdata-cellranger-vdj-GRCm38-alts-ensembl-7.0.0, which is based on the GRCm38 (Ensembl) genome assembly. Downstream analyses were performed in R v4.3.2 using Seurat v5.0.3 Ambient RNA contamination was corrected using SoupX(*62*). Quality control was performed separately for each age group, excluding low-quality cells based on the number of features, UMIs, and mitochondrial percent. Specifically, cells were retained if nFeature_RNA > 200 and percent.mt < 5%, while age-specific upper cut-offs were applied as follows: P0 (< 5,000 features, < 20,000 UMIs); P6 (< 5,000 features, < 21,000 UMIs); P10 (< 4,800 features, < 22,00 0 UMIs); P15 (< 5,000 features, < 22,000 UMIs); P21 (< 4,500 features, < 18,000 UMIs); Ad1 (< 5,000 feat ures, < 20,000 UMIs); Ad2 (< 5,000 features, < 20,000 UMIs). Doublets were removed using ScDblFinder(*63*) v1.16.0. Following quality control, datasets were merged and normalized using the SCTranform function with mitochondrial content regression, and integrated using Harmony(*64*). Dimensionality reduction was performed using principal component analysis (PCA; retaining the first 30 harmonized PCs), followed by Uniform Manifold Approximation and Projection (UMAP; dims = 1:30) for visualization.

Antibody-derived tag (ADT) matrices were then incorporated onto the single cell object for pre-processing using the dsb package(*65*) v1.0.3. Isotype control antibodies were used to estimate background noise and normalize protein expression using to account for technical noise and unspecific binding. Dimensionality reduction was conducted using PCA restricted to non-isotype markers, and the number of dimensions was determined as per above, using the elbow plot inflection point.

To perform the weighted nearest neighbor (WNN), variable features were identified separately for the RNA (SCT assay) and protein (ADT assay) modalities. Dimensionality reduction was applied using Harmony for RNA and PCA for ADT data. Clustering was performed on the weighted shared nearest neighbor (wsnn) graph using the Leiden algorithm at a resolution of 1. Singleton groups were retained to preserve rare populations. UMAP was run on the WNN graph to visualize the integrated data, using the computed weights and reduced dimensions. Cell identities from clustering were stored as metadata under `cell type` and `idents` for downstream analysis and visualization. To enable direct comparison of cell type proportions, each age group was downsampled to 3,500 cells. For the correlation heatmap, the relative abundance of each cluster was then calculated per age, and pairwise Pearson correlations of these distributions were then used to construct the heatmap. Ligand-receptor analysis was performed using LIANA(*66*).

#### B cell analysis

B cells were subclustered and reprocessed independently as per above from the RNA assay to investigate lineage-specific heterogeneity. Clustering was performed using the Leiden algorithm at a resolution of 1 using the WNN analysis. To explore transcriptional dynamics within B cells, PHATE v1.0.7 dimensionality reduction was performed(*66*) on library-size-normalized, square-root-transformed raw counts after PCA (n = 80PCs). Trajectory analysis was performed using Monocle3(*67*) using the WNN UMAP as dimensionality reduction. Pseudotime was computed using pro B cells as the root node to anchor the trajectory. BCR sequencing analysis was conducted using Platypus(*68*) v3.6.0 where clonal dynamics were analyzed using the VDJ_dynamics function, and V-J amino acid sequences were traced across development with P0 set as a reference repertoire.

#### MC data integration

Raw gene expression matrices were obtained from in-house CITE-seq experiment, and published single-cell RNA-seq datasets (Rustenhoven et al., 2021; Tauber et al., 2023). For Rustenhoven data, raw 10x Genomics CellRanger output was loaded using the Read10X function in Seurat (v4.3). For each dataset, a Seurat object was created using CreateSeuratObject with default settings filtering cells with <200 or >6,000 genes, >5% mitochondrial reads, or >30,000 UMIs. Following quality control, datasets were annotated with their respective sample origins and merged into a single Seurat object. To control for technical variation, we applied SCTransform normalization, performed independently on each dataset prior to integration. A set of 3000 highly variable genes was selected with SelectIntegrationFeatures. After merging, variable features were assigned across the combined object. Dimensionality reduction was performed via PCA followed by Harmony correction (retaining 30 dimensions). Cells were clustered by Leiden algorithm and mast cells were identified based on expression of Kit, Mrgprb2, and Cpa3. Mast-cell clusters from all datasets were subset and re-embedded. Gene-set scores for connective tissue and mucosal mast cells were calculated using Ucell(*69*).

#### Dani, et al. data reanalysis

Normalized counts and metadata table with cell annotations were used to reanalyze the single cell data from Dani, et al. Bar plots for immune cell percentages and violin plots were constructed as per above using the Seurat package.

#### Behavioral Assays

All behavioral testing was conducted in 8–16-week-old mice. Mice were habituated to the experimenter before testing by gently handling them for 10 min every day for a week. Mice were habituated to the testing room for a week before testing and for 1-2h prior to each assay. For each test, the apparatus and objects used were cleaned thoroughly with 70% ethanol between trials. Ethovision software was used for manual and scoring automatic quantification in all assays.

***Open field test*:** Mice were assessed in the open field chamber over a 10-minute period. Their activity was tracked using the Ethoviosion (Noldus) software to obtain the total distance traveled, the velocity, time spent freezing or in motion, as well as the time in the center zone of the field.

***Novel object recognition***: Novel object recognition was conducted as previously described. Mice were habituated for 10-minute sessions in the open field for three days. On the fourth day, mice were placed in the open field apparatus with two identical objects and allowed to explore for 10 min. The next day, the mice were tested, and a new object was introduced to the open field. “Interaction time” was scored manually when the nose of the test mouse interacted with the objects.

***Novel object location***: Novel object recognition was conducted as previously described. Mice were placed in the open field apparatus with two identical objects and allowed to explore for 10 min. The next day, the mice were tested, and one of the object’s placements was changed. “Interaction time” was scored manually when the nose of the test mouse interacted with the objects.

***Elevated plus maze*:** Mice were introduced to an elevated plus maze and assessed over 10 minutes. Zones were draws in the open and close arms of the maze and the Ethoviosion (Noldus) software was used to obtain the cumulative time spent in the open and closed arms.

***Social interaction test*:** Mice were habituated for 10 min to the three-chamber open field one day prior to testing. To test sociability, on test day 1, an adult age-matched mouse (female or male of 8-14 weeks) was placed in a wire cup in the left or right chamber (Social cup) and an object was placed under the wire cup was in the opposite chamber. The experimental mice were placed in the center chamber and allowed to explore the three-chamber open field for 10 minutes freely. Their exploration was recorded and tracked using the Ethoviosion software. The location of the social cup was alternated between the left or right chamber. Interaction was manually scored when the nose of the test mouse came in contact with one of the cups.

***Social novelty test:*** Twenty-four hours after the Social Interaction Test, mice were tested for social memory. The previously presented mouse from day 1 was placed in the same wire cup (Familiar cup) in the same chamber, and a novel, non-cage mate was placed in the opposite cup (Novel cup). The experimental mice were placed in the center chamber and allowed to freely explore the three-chamber open field for 10 minutes. Their exploration was recorded and tracked as in the social interaction test with manual scoring using Ethovision Software.

### QUANTIFICATION AND STATISTICAL ANALYSIS

Statistical analyses were performed using GraphPad Prism (v10.2.2, GraphPad Software), except for single-cell RNA sequencing analyses, which were conducted in R v4.3.2. For comparisons between two groups, normality was assessed using the Shapiro-Wilk test. Normally distributed data were analyzed using two-tailed Student’s t-tests; otherwise, the non-parametric Mann-Whitney test was applied. For comparisons involving more than two groups with a single independent variable, one-way ANOVA followed by Dunnett’s or Tukey’s post hoc tests was used when normality assumptions were met; otherwise, the Kruskal-Wallis test with Dunn’s multiple comparisons was applied. For analyses involving two independent variables, two-way ANOVA was performed, followed by Tukey’s, Bonferroni’s, or Sidak’s post hoc tests, as specified in figure legends. Data are presented as mean ± standard error of the mean (SEM). p < 0.05 was considered significant: *p < 0.05; **p < 0.01; ***p < 0.001; ****p < 0.0001. p > 0.05 was considered nonsignificant (ns). Sample sizes, number of independent experiments, and sex of animals are detailed in the respective figure legends.

## Acknowledgments

We thank all members of the Deczkowska lab, past and present, for technical support and fruitful discussions. We are grateful to the Central Animal Facility of Institut Pasteur for taking care of the animals used in this study, the C2RT, especially the Flow Cytometry, Photonic BioImaging (PBI), Single Cell Biomarkers, Biomics, and Mass Spectormetry for biology (MSBio) platforms for their technical support. We thank A. Blondeau for technical support and editing. We thank A. Patrizi and E. Gomez Perdiguero for fruitful discussions.

## Funding

The research in Deczkowska lab is supported by the G5 funding from Institut Pasteur, the European Research Council Starting grant (BrainGate), ERA-Net Neuron 2024 (BodyMood), Agence Nationale de la Recherche (DevoCP), Ville de Paris EMERGENCE(S) grant, Alzheimer’s Association Research Grant (AARG-22-917964), Fédération pour la Recherche sur le Cerveau, ERA-Net NEURON 2024 and Don Explore AD (Programme Explore de l’Institut Pasteur). Our lab is part of the DIM C-BRAINS, funded by the Conseil Régional d’Île-de-France. This research was funded, in whole or in part, by French National Research Agency (DevoCP) and ERC (BrainGate). For the purpose of open access, the author has applied a CC- BY public copyright license to any Author Manuscript version arising from this submission.

## Author contribution

Conceptualization and methodology: SAM, LT, and AD. Investigation: SAM, IB, TM, ABM, MK, ARP, XD, NG, FJ, and LT. Data analysis: SAM, LT.

Bioinformatics analysis: SAM, JJ, and OM. Human CP imaging: CT, and MH. Proteomics data analysis: VP, and MM. Writing—original draft: SAM, LT, and AD.

Writing—review & editing: SAM, LT, and AD. Funding acquisition: AD.

Resources: AD. Supervision: LT, and AD.

## Competing interests

The authors declare no competing interests.

## Data and materials availability

Single-cell RNA-seq data have been deposited at GEO with the accession code GSE310655. CSF proteomics data have been deposited at PRIDE with the accession code PXD071204. Further information and requests for resources and reagents should be directed to and will be fulfilled by the lead contact, Aleksandra Deczkowska.

---

**Fig. S1.**
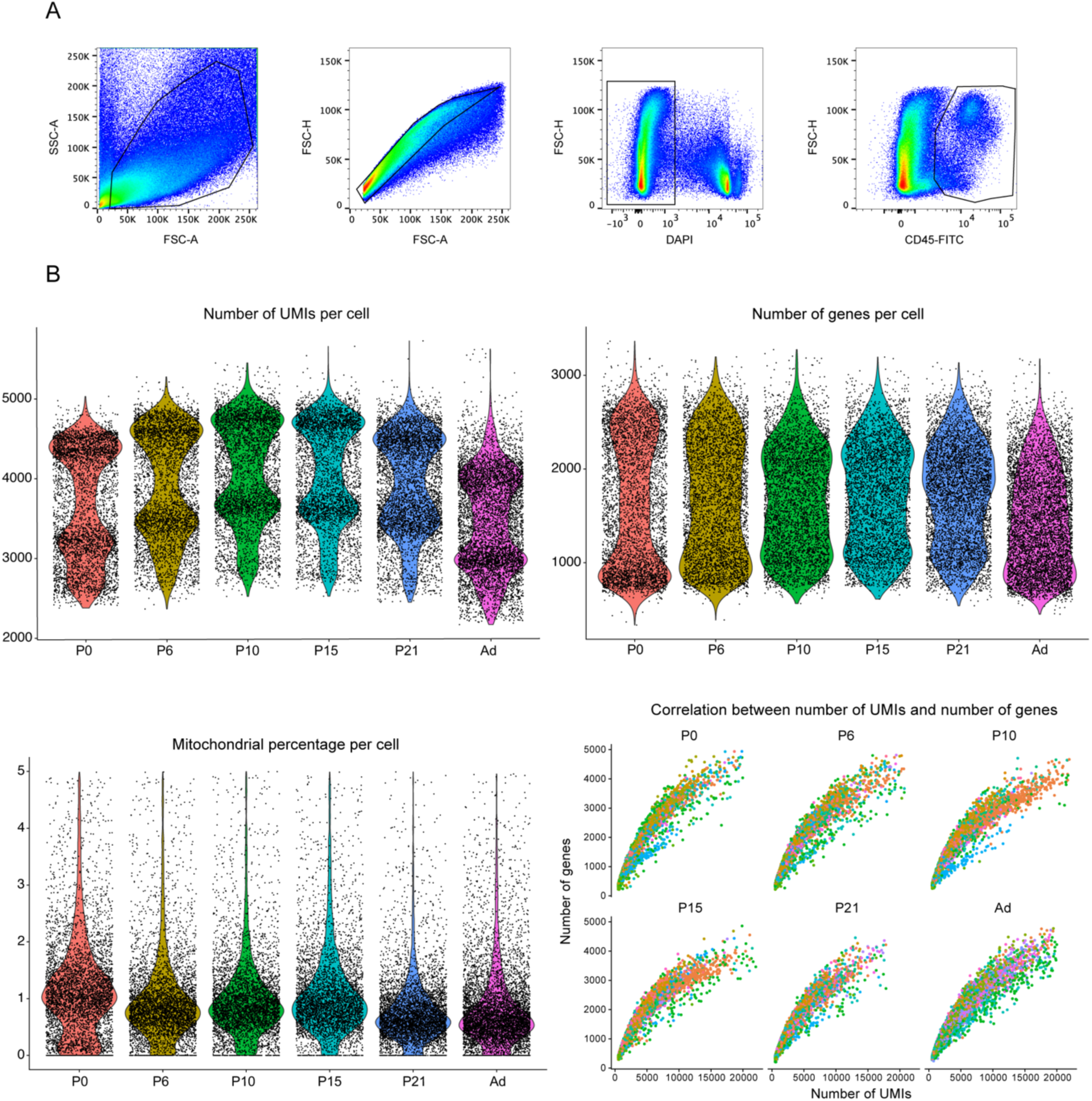
Quality control metrics for CITEseq. **(A)** Gating strategy for fluorescence activated cell sorting of CP immune cells for CITE-seq in Fig.1. **(B)** Quality control measures for each sample post filtering: percentage of mitochondrial genes, number of genes, number of UMIs captured and correlation between number of genes and UMIs.

**Fig. S2.**
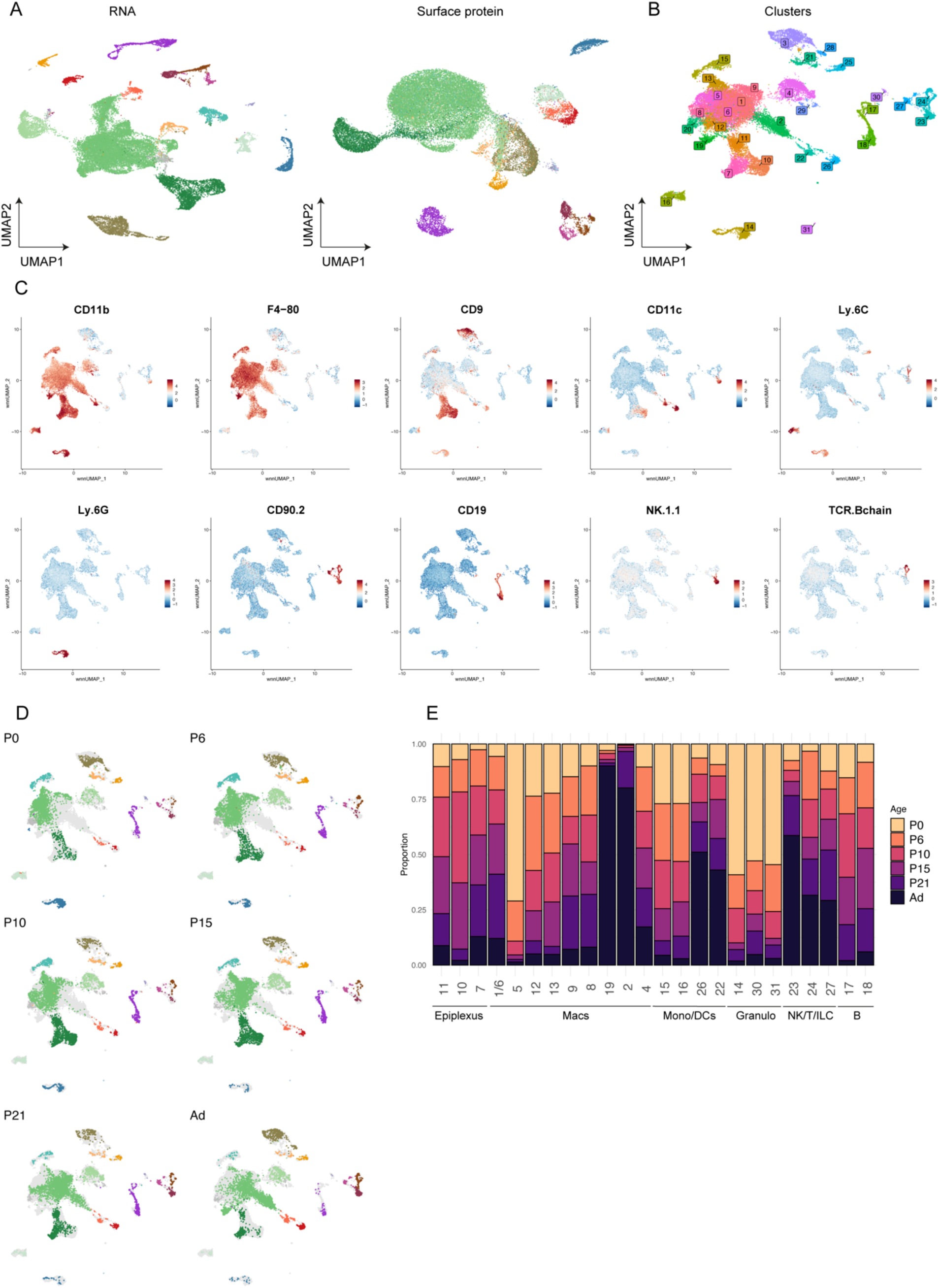
Mapping immune cell dynamics during postnatal development in the CP. **(A)** UMAP visualization of RNA (right) and surface protein (left) based dimensionality reduction. **(B)** Integrated UMAP visualization of all 34 clusters obtained using the WNN analysis. **(C)** Featureplots showing normalized expression of marker surface proteins: CD11b: myeloid cells; F4/80: monocytes/macrophages; CD9: epiplexus cells; CD11c: dendritic cells; Ly6C: monocytes; Ly6G: neutrophils; CD90.2: lymphocytes; CD19: B cells; NK1.1: NK cells; TCRbeta: T cells. **(D)** Representative UMAPs separated per age. Grey background represents the integrated UMAP across all ages. **(E)** Bar plots showing the proportion of individual immune clusters during postnatal development (subsampled on 3000 immune cells per age).

**Fig. S3.**
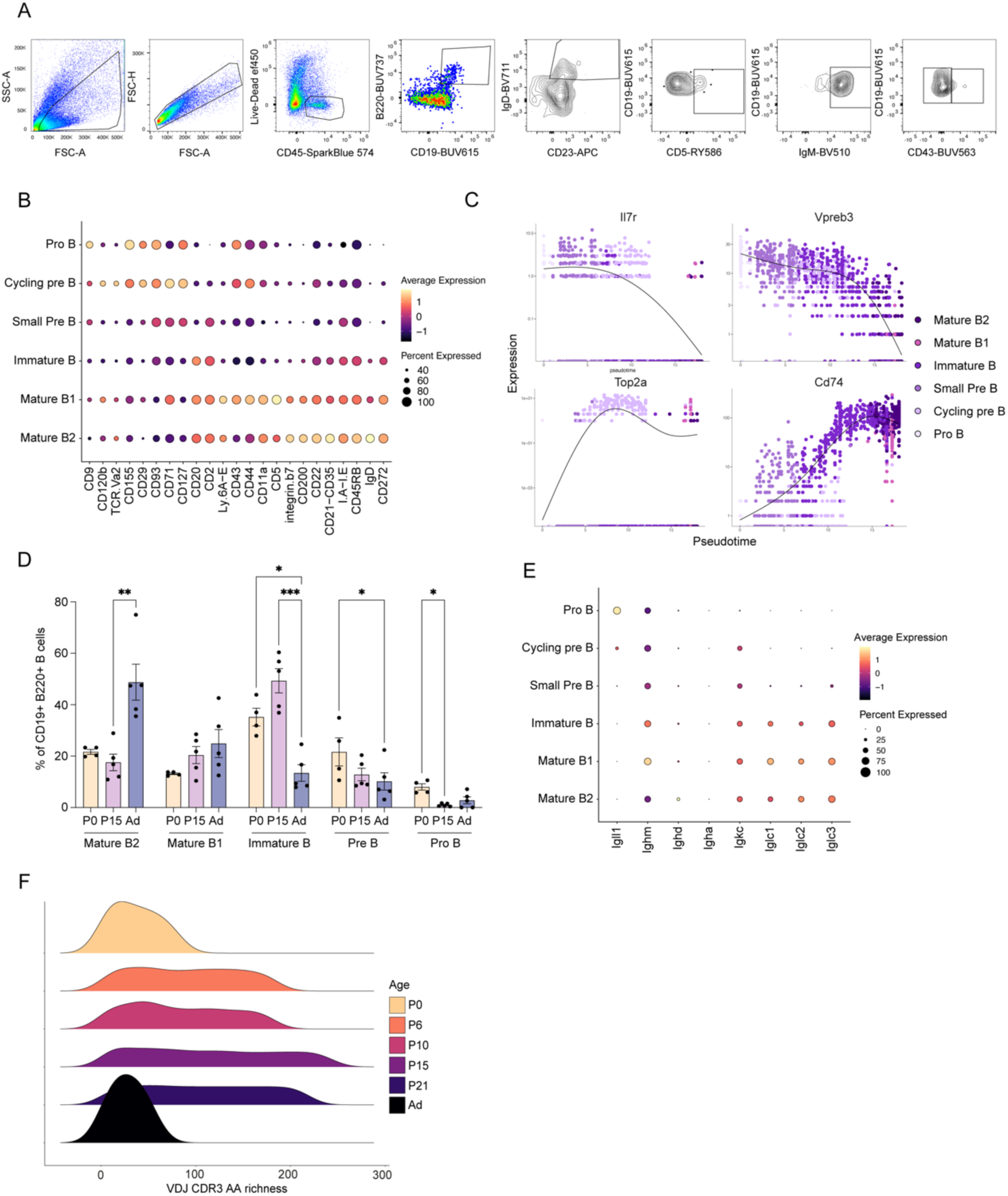
The CP is a developmental niche for B cell development. **(A)** Gating strategy for flow cytometric identification of B cell subpopulations. **(B)** Dot plot showing normalized expression of top differential marker proteins for the identified B cell subpopulations. The color represents mean expression level, and the size indicates the proportions of cells expressing the proteins. **(C)** Representative differentially expressed genes exhibiting temporally distinct expression dynamics along pseudotime, identified using Monocle3. **(D)** Percentage of B cell subpopulations in the developing CP identified by flow cytometry. n= 4-5 mice per group. Data are represented as mean ± SEM, one-way ANOVA with Tukey’s multiple comparison test. **(E)** Dotplot showing immunoglobulin gene expression in B cells. **(F)** BCR repertoire diversity across CDR3 sequence lengths and age.

**Fig. S4.**
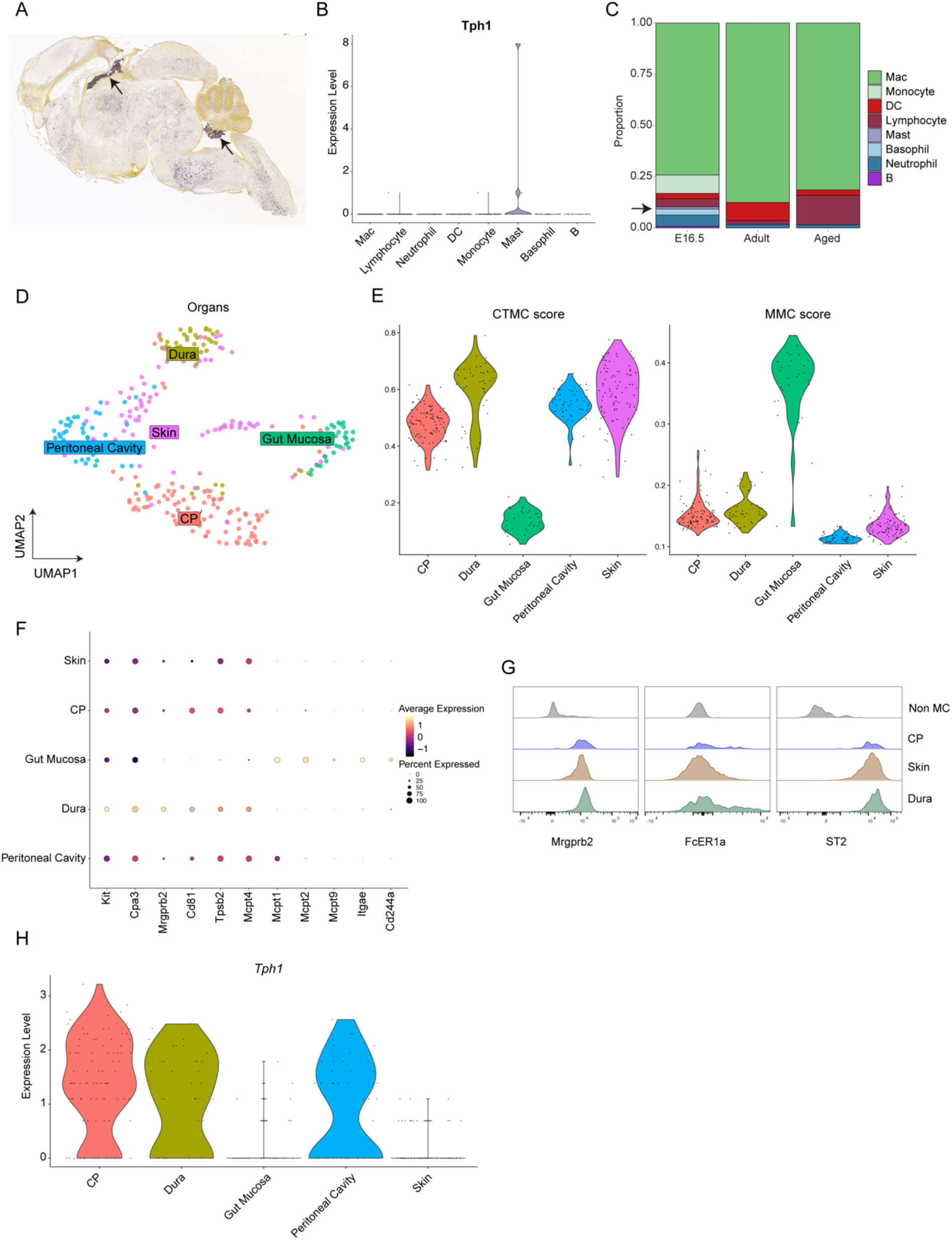
Characterization of CPMCs. **(A)** Expression of *Htr2c* in the P4 mouse brain. Allen Developing Mouse Brain Atlas, developingmouse.brain-map.org/experiment/show/100047619. **(B)** Bar plots showing the immune composition of the CP in from embryonic development to aging from Dani, et al. 2021(*19*). Arrow indicates MCs. **(C)** Violin plots depicting the expression of *Tph1* across immune cells. Data reanalyzed from Dani, et al. 2021(*19*). **(D)** Integrated UMAP of MCs distribution across tissues. Data integrated from Tauber et al(*45*) and Rustenhoven et al(*17*). **(E)** Violin plot showing the normalized CTMC and MMC score expression based on key marker genes; (see marker genes in Table S1). **(F)** Dot plot of marker genes for CTMCs and MMCs across organs **(G)** Representative flow histograms depicting protein expression of phenotypic markers from neonatal MrgprB2-EGFP/Cre mice across various organs. **(H)** Violin plot depicting *Tph1* expression across organs.

**Fig. S5.**
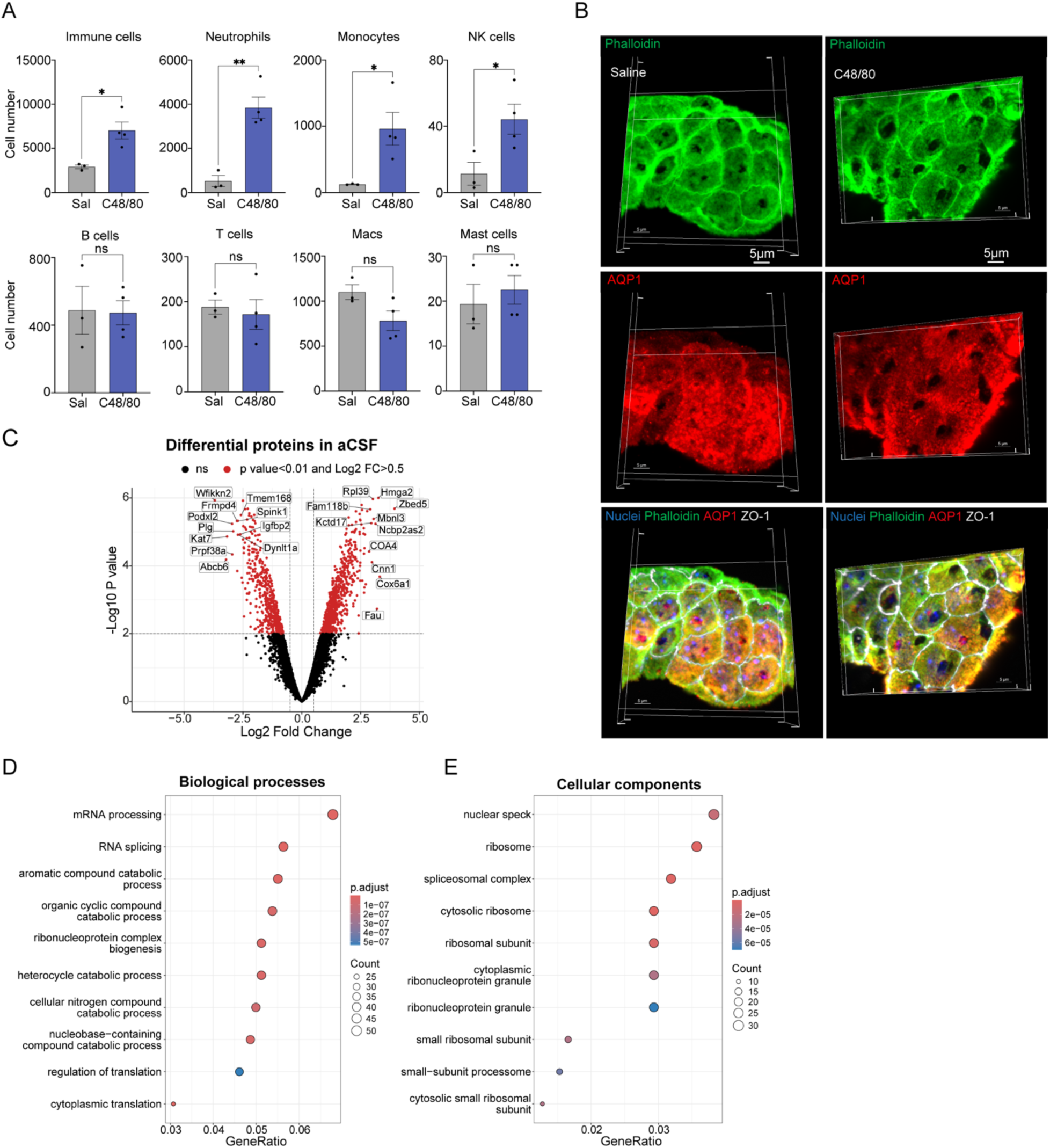
MC activation induces apocrine secretion. **(A)** Flow cytometry quantification of total number of immune cells, neutrophils, monocytes, NK cells, T cells, B cells, Macs, and MCs in the CP 24h post i.c.v injection of C48/80. Data are represented as mean ± SEM. P values were calculated by unpaired t-test. **(B)** Representative three-dimensional reconstruction of confocal imaging of 4V CP sections stained for nuclei, phalloidin, AQP1 and ZO-1, three hours following i.c.v injection of saline or C48/80. Bar: 5µm. **(C)** Volcano plot of the differentially expressed proteins in the aCSF of C48/80 stimulated and control CP explants. **(D)** Dot plot showing top10 statistically most significant BP terms enriched in the C48/80 aCSF proteome. **(E)** Dot plot showing top10 statistically most significant CC terms enriched in the C48/80 aCSF proteome. *p < 0.05; **p < 0.01; ns: p > 0.05.

**Fig. S6.**
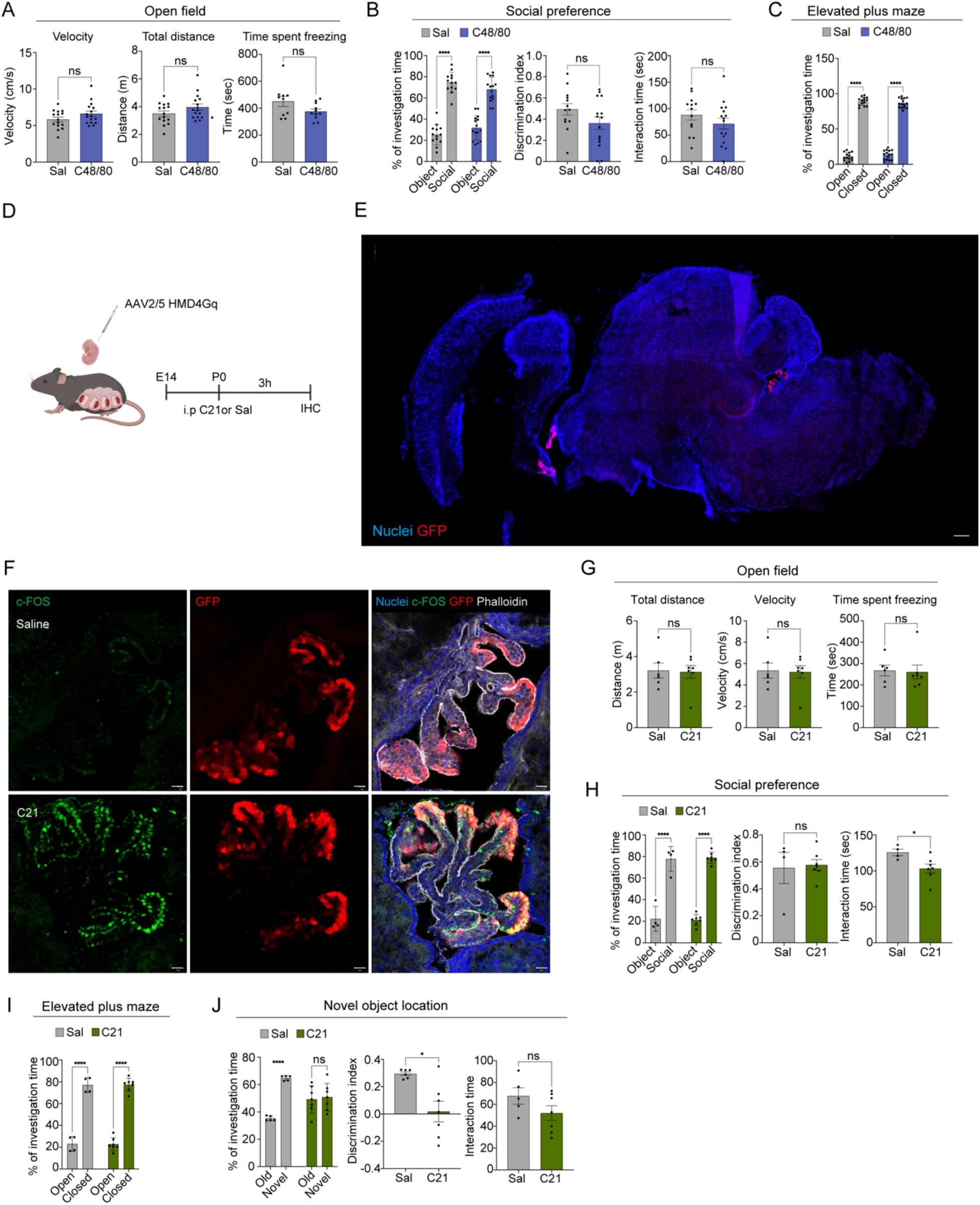
MC activation and CPEC activation in neonates does not impact activity, social preference, or anxiety. **(A) Left**: Velocity, **Center**: Total distance moved, **Right**: total time spent freezing in the open field test by control and C48/80 treated mice. Data were analyzed by Mann-Whitney test. **(B) Left**: Percentage of interaction time spent with the social versus object cup control and C48/80 injected mice in the social preference test. Data analyzed by two-way ANOVA with Tukey’s multiple comparisons test. **Center**: Discrimination index for control and C48/80 mice. **Right**: total interaction time during the social preference test by control and C48/80 mice. Data were analyzed by Mann-Whitney test. **(C)** Percent time spent in closed vs. open arms by controls and C48/80 mice. Two-way ANOVA with Tukey’s multiple comparisons test. **(D)** Experimental scheme for chemogenetic manipulation of CPECs. **(E)** Representative confocal image depicting DREADD expression in the P0 brain upon AAV2/5 delivery at E14. **(F)** Representative confocal images of the 4V CPEC activation of P0 mice three hours following injection with C21 or saline. **(G) Left**: Total distance moved, **Center**: velocity, **Right**: total time spent freezing in the open field test by control and C21-treated mice. Data were analyzed by Mann-Whitney test. **(H) Left**: Percentage of interaction time spent with the social versus object cup control and C21-injected mice in the social preference test. Data analyzed by two-way ANOVA with Tukey’s multiple comparisons test. **Center**: Discrimination index for control and C21 mice. **Right**: total interaction time during the social preference test by control and C21 mice. Data were analyzed by Mann-Whitney test. **(I)** Percent time spent in closed vs. open arms by controls and C21 mice. Two-way ANOVA with Tukey’s multiple comparisons test. **(J) Left**: Percentage of interaction time spent with the novel versus old location for the object in saline and C21injected DREADD mice. Data were analyzed by two-way ANOVA with Tukey’s multiple comparisons test. **Center**: Discrimination index for saline and C2-1injected DREADD mice. Data were analyzed by Mann-Whitney test. **Right**: total interaction time with the novel and old object location by saline C21-injected DREADD mice. Data were analyzed by Mann-Whitney test. *p < 0.05; **p < 0.01; ***p < 0.001; ****p < 0.0001. ns: p > 0.05.

Table S1. Gene list used for CTMC and MMC score generation (from Tauber et. al (45)).

**Table S2.** Materials and reagents.

| **REAGENT or RESOURCE** | **SOURCE** | **IDENTIFIER** |
| --- | --- | --- |
| **Antibodies** |  |  |
| TotalSeq Cmouse universal cocktail V1 | Biolegend | Cat# 199903 |
| TruStain FcX plus | Biolegend | Cat# 156603 |
| Fc Block CD16/32 | BD Biosciences | Cat# 553141 |
| CD45-FITC | Biolegend | Cat# 103108 |
| CD43-BUV563 | BD Biosciences | Cat# 741238 |
| CD19-BUV615 | BD Biosciences | Cat# 751213 |
| B220-BUV737 | BD Biosciences | Cat# 612838 |
| CXCR4-BV421 | Biolegend | Cat# 146511 |
| IgM-BV510 | Biolegend | Cat # 406531 |
| IgD-BV711 | Biolegend | Cat # 405731 |
| CD45-Spark-Blue574 | Biolegend | Cat# 103184 |
| CD5-RY585 | BD Biosciences | Cat# 753291 |
| CD11b-PE/DAZZLE594 | Biolegend | Cat# 101256 |
| CD24-PE/Cy7 | Biolegend | Cat# 101821 |
| CD23-APC | Biolegend | Cat# 101619 |
| CXCR5-APC/Cy7 | Biolegend | Cat# 145525 |
| FcER1a-BUV805 | BD Biosciences | Cat # 749151 |
| c-KIT-RB744 | BD Biosciences | Cat# 570556 |
| Lin-PE | Biolegend | Cat # 133303 |
| CD200R-APC | Biolegend | Cat# 123916 |
| ST2-BB700 | BD Biosciences | Cat # 746115 |
| CD14-BUV395 | BD Biosciences | Cat # 740241 |
| NK1.1-BUV805 | BD Biosciences | Cat# 569164 |
| B220-BV421 | BD Biosciences | Cat # 562922 |
| CX3CR1-BV570 | Biolegend | Cat# 149061 |
| Ly6C-BV711 | Biolegend | Cat # 128037 |
| TCRb-PE/Cy7 | Biolegend | Cat# 109221 |
| Ly6G-AF647 | Biolegend | Cat# 127609 |
| F4/80-APC/Fire810 | Biolegend | Cat# 123165 |
| Mouse IgG1 anti mouse ZO-1 (clone 1A12) | Invitrogen | Cat# 33-9100 |
| Rat anti mouse CD31 | BD Pharmingen | Cat# 550274 |
| Rat IgG2a anti mouse CD45R/B220 - AF647 | Biolegend | Cat# 103229 |
| Rat IgG2a anti Serotonin | Novus Biologicals | Cat# NB100-65037 |
| Rabbit anti mouse AQP1 | Merck | Cat# AB2219 |
| Chicken anti GFP | Abcam | Cat# ab13970 |
| Rabbit anti mouse c-FOS | Abcam | Cat# ab190289 |
| Goat anti mouse IgG1 AF546 | Invitrogen | Cat# A21123 |
| Donkey anti rat DyLight 488 | Thermo Fisher Scientific | Cat# SA5-10026 |
| Donkey anti rat DyLight 550 | Thermo Fisher Scientific | Cat# SA5-10027 |
| Donkey anti rabbit DyLight 550 | Thermo Fisher Scientific | Cat# SA5-10039 |
| Goat anti chicken AF488 | Invitrogen | Cat# A11039 |
| Goat anti mouse IgG1 AF647 | Invitrogen | Cat# A21240 |
| **Chemicals, peptides, and recombinant proteins** |  |  |
| Phalloidin-AF488 | Cell Signaling Technology | Cat# 8878 |
| Phalloidin-AF660 | Invitrogen | Cat# A22285 |
| Avidin-FITC | BioLegend | Cat# 405102 |
| Live-dead fixable violet deadkits | Thermo Fisher Scientific | Cat# 10645203 |
| Compound 48/80 | Sigma | Cat# C2313-100MG |
| DAPI | Thermo Fisher Scientific | Cat# 10116287 |
| SB 242084 Hydrochloride | Sigma | Cat# 5064170001 |
| Actinomycin D | Thermo Fisher Scientific | Cat# 10247270 |
| Compound 21 | Bio techne sas | Cat# 5548/10 |
| Collagenase IV | Thermo Fisher Scientific | Cat# 10780004 |
| Collagenase | Thermo Fisher Scientific | Cat# 11500536 |
| DNASE 1 | New England Biolabs | Cat# M0303S |
| Albumin | Thermo Fisher Scientific | Cat# 11410262 |
| **Commercial kits** |  |  |
| NextSeq 1000/2000 P2 reagents | Illumina | Cat# 20046811 |
| Chromium next GEM single cell | 10X Genomics | Cat# 1000265 |
| library construction kit | 10X Genomics | Cat# 1000250 |
| dual index kit TN set | 10X Genomics | Cat# 1000254 |
| chromium next GEM chip K single cell kit | 10X Genomics | Cat#1000190 |
| chromium single cell mouse BCR amplification | 10X Genomics | Cat# 1000255 |
| dual index kit TT set | 10X Genomics | Cat# 1000287 |
| 5' feature barcode kit | 10X Genomics | Cat# 1000541 |
| **Experimental models:**<br>**Organisms/strains** |  |  |
| Mouse: C57BL/6J | Janvierlabs |  |
| Mrgprb2-Cre/GFP | Provided by X.Dong and N.Gaudenzio | https://www.nature.com/articles/nature14022 |
| **Virus strains** |  |  |
| AAV2/5-CMV>hM3DGq-EGFP | VectorBuilder | Manufactured on demand |
| **Deposited data** |  |  |
| CITEseq CP P0-Ad |  | GSE310655 |
| CSF and aCSF proteomics |  | PRIDE database: PXD071204 |
| **Software and algorithms** |  |  |
| Seurat v5.0.3 |  |  |
| R v4.3.2 |  |  |
| dsb v1.0.3 |  |  |
| Ucell v2.7.3 |  |  |
| LIANA v0.1.13 |  |  |
| PHATE v1.0.7 |  |  |
| Platypus v3.6.0 |  |  |
| ClusterProfiler v4.10.1 |  |  |
| Ethovision (Noldus) |  |  |
| GraphPad Prism v10.2.2 |  |  |

## Notes

### Competing Interest Statement

The authors have declared no competing interest.

